# Structure-function relationships in mitochondrial transcriptional condensates

**DOI:** 10.1101/2021.12.30.474545

**Authors:** Marina Feric, Azadeh Sarfallah, Furqan Dar, Dmitry Temiakov, Rohit V. Pappu, Tom Misteli

## Abstract

Phase separation organizes many membraneless structures in cells. The functional consequences of concentrating cellular machinery into biomolecular condensates, however, are largely unclear. One fundamental cellular function that has been linked to condensate formation is transcription. Here, we have reconstituted mitochondrial transcription in condensates from purified components. We find that the core components of the mt-transcriptional machinery form multi-phasic, viscoelastic condensates *in vitro*. Strikingly, the rates of condensate-mediated transcription are substantially lower than equivalent reactions in bulk solution. These condensate-mediated decreases in transcriptional rates are associated with the formation of dynamically arrested vesicular structures that are driven by the production and accumulation of RNA during transcription. Using coarse-grained, equilibrium simulations, we show that the generation of RNA alters the phase behavior and the organization of transcriptional components within condensates and that the *in vitro* mtcondensates are non-equilibrium structures. Together, our *in vitro* and *in silico* approaches shed light on how proteins and (ribo)nucleic acids biophysically self-assemble within mitochondria *in vivo*. Our results highlight the complex morphologies of transcribing, multicomponent condensates and they illustrate the interdependent structure-function relationships in condensates.

**Significance Statement:** Mitochondria condense their genome into transcriptionally active mt-nucleoids. These structures fit the definition of biomolecular condensates that form via macromolecular phase separation. We take advantage of the ability to reconstitute mitochondrial transcriptional condensates in vitro from minimal components. We find that the production and accumulation of RNA alters the phase behavior of transcriptional condensates. The altered phase behavior is linked to the formation of arrested, non-equilibrium vesicular structures. Similar changes to phase behavior of proteins and (ribo)nucleic acids can be recapitulated in live mitochondria through knockdown of mt-nucleoid core components. Computer simulations help identify biophysical mechanisms that are needed to maintain the steady-state structures of transcriptional condensates.

## Introduction

Proteins and nucleic acids form diverse biomolecular condensates that are proposed to arise via macromolecular phase separation (1-4). Condensates emerge by demixing of protein, RNA and DNA components from their cellular surroundings to form distinct, non-membrane bound cellular structures. The formation of condensates is typically mediated by multivalent homotypic and heterotypic interactions amongst proteins and nucleic acids (1). Prominent biomolecular condensates include P-granules and stress granules in the cytoplasm (5, 6) as well as the nucleolus and RNA splicing factor speckles in the nucleus (7, 8). Within mitochondria, the genome-containing mitochondrial (mt)-nucleoid and RNA processing granules are condensates that appear to form via phase separation (9, 10).

Condensates are enriched in functional components, such as transcription factors or RNA processing factors. Increased concentrations of bioactive macromolecules within condensates is thought to be important for enhancing the rates of key biochemical reactions within condensates (11). However, the connection between condensate structure and function remains unclear (2). A major hurdle in elucidating structure-function relationships of condensates has been the difficulty to reconstitute functionally active condensates *in vitro* with all the biochemically relevant components. Conversely, *in vivo* perturbations, such as mutagenesis or pharmacological disruption, make it challenging to directly isolate the effects of phase behavior from the functional properties of the affected cellular components.

Condensates are thought to contribute to many cellular functions, including genome organization and transcription (12-14). Major architectural chromatin proteins such as the linker histone H1 (15) and the heterochromatin protein HP1*α* form condensates *in vitro* and *in vivo* (16, 17). Phase separation, in different manifestations, has been suggested to contribute to higher-order organization of genomes into domains and compartments (18-20). In particular, various components of the transcription machinery spontaneously concentrate into condensed phases in the mammalian nucleus, including prominently at sites of super-enhancers (21). This behavior has been attributed to the intrinsically disordered regions (IDRs) found in many transcription factors and chromatin proteins (22). IDRs are thought to mediate weak, multivalent protein-protein interactions that give rise to dynamic, non-stoichiometric condensed assemblies (23). Functionally, the condensation of transcription components is associated with bursts of transcription of RNA (24) that are consequently followed by dissolution of the condensate in an effective feedback loop (25).

The mitochondrial genome (mtDNA) and its own dedicated gene expression machinery are also organized via phase separation (9). Human mitochondria contains hundreds of copies of their own 16 kb, circular genome (26) that assemble into mitochondrial nucleoids, which are membraneless, nucleoprotein complexes of ∼100 nm in diameter containing mtDNA and associated proteins (27, 28). In support of phase separation as a driver of mt-nucleoid organization, the major mt-genome architectural protein TFAM phase separates *in vitro* and *in vivo*, and combined with mtDNA, forms condensates that recapitulate the behavior of mt-nucleoids in cells (9). The mt-nucleoids serve as sites of transcription of long, polycistronic mtRNA, which becomes further processed in adjacent RNA granules that associate with the mitochondrial membrane and are also thought to form via phase separation (10).

The relative simplicity, involving only a small number of minimally required components, makes mitochondrial transcription a unique and tractable model system to probe structure-function relationships in a biologically relevant condensate. Mitochondrial transcription can be reconstituted under soluble conditions with only four components: mtDNA, the single-subunit mitochondrial RNA polymerase POLRMT, and two transcription factors, TFAM and TFB2M (29, 30). Here, we have reconstituted mitochondrial transcription under condensate-forming conditions *in vitro*, and we directly probe functional consequences of the condensate morphologies and their internal physicochemical environments. We demonstrate that the mitochondrial transcription machinery forms multi-phasic, non-equilibrium condensates with slow diffusivities, contributing to dampened transcriptional kinetics compared to equivalent reactions in bulk solutions. Importantly, we find that the production of nascent RNA during transcription alters the structure of the condensate. Our results demonstrate a close interplay between the physical behavior and functional activity of an archetypal biomolecular condensate.

## Results

### Individual components of the mitochondrial transcription machinery undergo phase separation *in vitro*

Mitochondrial transcription has previously been reconstituted with three proteins and a DNA template containing a mitochondrial promoter under dilute conditions (30). Given the organization of mt-nucleoids into condensates within the crowded mitochondrial matrix *in vivo* (9), we sought to establish conditions for *in vitro* mitochondrial transcription in reconstituted condensates.

Taking a bottom-up approach, we first established the individual phase behavior of the minimal components required for mitochondrial transcription. The transcription factors TFAM and TFB2M combined with the polymerase POLRMT represent the minimal components of human mitochondrial transcriptional machinery. Structural studies (31) and bioinformatics analysis show that these proteins contain a combination of ordered domains and IDRs (Fig. 1A). Computational predictions for disordered proteins suggest that unbound TFAM is the most disordered of the three proteins with a flexible linker that bridges two DNA binding domains (High Mobility Groups A, B) and a disordered C-terminus. These features are consistent with the flexible nature and multiple conformations that have been reported for TFAM molecules in solution (32). TFB2M and POLRMT are significantly more structured with well-folded, functional domains (29, 33), but both contain disordered regions at their N-termini. The modular nature of these proteins suggests that they have the potential to drive or contribute to phase separation of mt-nucleoids, as disordered regions contribute to non-specific, weak interactions and ordered regions provide specific, strong interactions (34).

**Figure 1:**
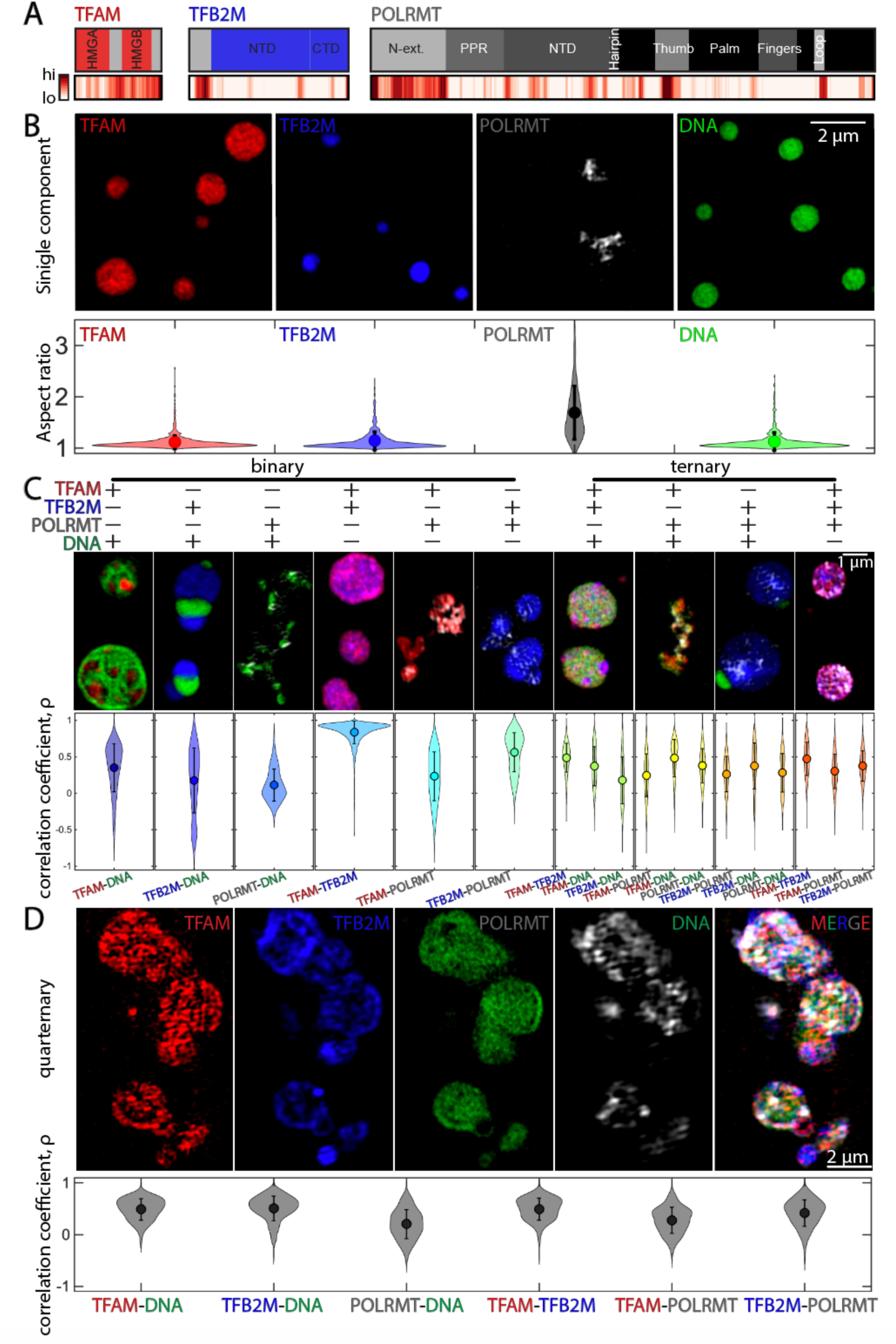
Equilibrium phase behavior of individual, binary, ternary, and quaternary condensates comprised of mt-transcriptional components. (A) Protein domain analysis for core mitochondrial transcription proteins TFAM, TFB2M, and POLRMT. Top row illustrates the known protein domains (red, blue, dark grey, respectively) and the unfolded, intrinsically disordered domains are indicated in light gray. TFAM contains two High Mobility Group domains (HMGA/B) separated by a disordered linker and flanked by a disordered C-tail. TFB2M contains an N-terminal domain (NTD) which adopts a methyltransferase fold, and the C-terminal domain (CTD), which consists of four alpha helices and a short, flexible “tail.” POLRMT is a single-subunit DNA-dependent RNA polymerase, which is distantly related to the bacteriophage T7 RNAP. It contains a hand shaped CTD that harbors the catalytic site POLRMT also contains an N-terminal domain attached to the catalytic core via a poly-proline linker. Bottom row shows probability of intrinsically disordered sequences as predicted by several models using D^2^P^2^, where high and low likelihoods for disorder are indicated in red (high) and white (low). (B) SIM images of condensates formed for individual components at room temperature on pluronic-treated coverslips: 10 µM TFAM in 10% PEG (red), 10 µM TFB2M in 10% PEG (blue), 1.5 µM POLRMT in 5% PEG (gray), and 500 nM DNA in 10% PEG (green). Bottom row contains violin plots of the aspect ratio for all condensates analyzed. n = 4 experimental replicates, average (dot) values are indicted, error bar represents the standard deviation. Scale bar = 2 µm. (C) Binary and ternary compound droplets. Violin plots of correlation coefficient measured for each pair of channels; correlation coefficient = 1 denotes complete colocalization. Scale bar = 1 µm. n = 4 experimental replicates, average (dot) values are indicted, and error bar represents the standard deviation. (See Table 1 for concentrations). (D) Quaternary droplets at room temperature. Top row includes individual channels of ∼6 µM TFAM (red), ∼6 µM TFB2M (blue), ∼6 µM POLRMT (gray), and ∼500 nM DNA (green), and the merged image in 5% PEG. Scale bar = 2 um. Bottom row contains violin plots of the correlation coefficient for all pairs of channels. n = 3 experimental replicates, average (dot) values are indicted, and error bar represents the standard deviation. Buffer for all conditions was 20 mM Tris-HCl, pH ∼8.0, 20 mM BME, 10 mM MgCl_2_, and ∼100 mM NaCl at room temperature.

In terms of phase behavior, we find that TFAM, TFB2M, and POLRMT as well as DNA individually demix from solution and form dense phases in the presence of the macromolecular crowder, polyethylene glycol (PEG, MW ∼3 kDa). Conditions that promote phase separation include 10 µM TFAM or TFB2M in 10% PEG; 1.5-10 µM POLRMT in 5-10% PEG; 500 nM DNA in 10% PEG) (Fig. 1B; Fig. S1). Condensates formed by TFAM, TFB2M, or DNA resemble highly spherical droplets with an aspect ratio of ∼1 (Fig. 1B; Fig. S1). In contrast, POLRMT assembles into highly irregular structures (Fig. 1B; Fig. S1). The differential behavior of POLRMT can be accounted for by the fact that phase separation is a density transition (35), whereby dense phases, concentrated in specific types of macromolecules, coexist with dilute phases. Density transitions can generate a spectrum of morphologies and dynamics for dense phases, especially for condensates formed by nucleic acids; moreover, the interplay between phase separation (the density transition) and percolation or gelation, which refers to the networking of macromolecules via physical crosslinks, often results in dynamically arrested phases (35-39). Accordingly, irregular mesoscale structures are more likely to imply the formation of dynamically arrested phases, representing metastable, non-equilibrium structures, wherein one or more macromolecules are immobile because they are a part of highly crosslinked networks. In line with this interpretation, the structures formed by POLRMT show limited recoveries after photobleaching (Fig. S1D,E) and fit the description of such dynamically arrested phases (39, 40). Overall, our results show that all minimal components of the mitochondrial transcription machinery can undergo phase separation via homotypic interactions, yet each forms condensates with distinct dynamics and/or morphologies.

### Multicomponent systems form heterogeneously organized and dynamically arrested condensates *in vitro*

Next, we documented the joint phase behavior of multiple components by characterizing the structures formed by binary and ternary mixtures of components of the minimal mitochondrial transcription machinery (Fig. 1C). Using the correlation coefficient as a metric of their colocalization, we find that TFAM and DNA form multi-phase condensates, containing micron-sized sub-domains that are either TFAM-rich or DNA-rich (Fig. 1C). In contrast, TFB2M and POLRMT show lower degrees of colocalization with DNA (Fig. 1C). When the proteins were mixed in pairs without DNA, TFAM and TFB2M colocalize with one another although they do not colocalize as well as with POLRMT (Fig. 1C). This again, is a likely consequence of POLRMT driving the formation of dynamically arrested phases.

For ternary combinations of pairs of proteins with DNA, the droplet organization remained inhomogeneous for all combinations, with micron-sized domains forming for TFAM-TFB2MDNA and TFB2M-POLRMT-DNA (Fig. 1C). However, for the ternary mixture of all proteins without DNA, the condensates became more well-mixed, implying that the heterotypic interactions of proteins with DNA significantly contribute to the emergence of spatially distinct coexisting phases of the compound droplets (Fig. 1C). These results demonstrate differential phase behavior of the various components of the mitochondrial transcription machinery.

Towards building transcriptionally competent condensates, all four biomolecular components (DNA, POLRMT, TFAM and TFB2M) were combined in equimolar protein ratios (∼6 µM TFAM, TFB2M, and POLRMT with 500 nM DNA) in the presence of 5% PEG (Fig. 1D). In this mixture all biomolecules condense into single droplets (Fig. 1D). However, these droplets do not fuse with neighboring droplets. Instead, they come into contact and form dynamically arrested, higher-order structures (Fig. 1D). This is reminiscent of arrested coalescence observed in non-biological soft matter (41). While interfacial tension should enable fusion and a lowering of the interfacial energy, the internal network structure of the droplet provides elastic resistance to droplet deformation and fusion. Coalescence can begin, but becomes arrested when the two energies are balanced at an intermediate stage of coalescence. This arrest can happen at different stages of coalescence and is tied to the interplay of the surface energy and the elastic energy associated with the internal network structure of droplets. The extent of balancing of interfacial tension and elastic energies determines the extent of metastability and the lifetimes of the arrested phases.

We also found that localization of individual components within the multi-phasic structures remained heterogeneous: POLRMT showed the lowest level of colocalization with all other components, while TFB2M tended to accumulate more peripherally (Fig. 1D). Importantly, the localization correlation coefficients for specific pairs tended to be higher in the compound droplet than those found in the binary or ternary droplets (Fig. 1C-D). This behavior is consistent with the dual-nature of TFAM, which has affinity for DNA via its N-terminal DNA-binding domain (HMGA), but also for other proteins, such as POLRMT, via its disordered C-terminus (9). Together, these results show that the minimal mitochondrial transcription machinery can collectively form multi-phasic, dynamically arrested condensates *in vitro*.

### Condensates support transcription and dampen transcription rates

Next, we sought to test whether transcription could be reconstituted *in vitro* under condensate-forming conditions. In pilot experiments, we first characterized the effect of increasing concentrations of reaction components starting with standard, soluble *in vitro* mt-transcription reactions (30, 42). We added a full set of nucleotides (NTPs) to the mixture of 0.6 µM TFAM, 0.6 µM TFB2M, 0.6 µM POLRMT, and 50 nM DNA in the absence of any crowder (–PEG), representing soluble conditions. After 30 minutes of incubation ∼35ºC (see SI Appendix), transcriptional activity was measured by detection of a ∼300 nt RNA product using a PCR-amplified template containing the LSP promoter and radioactively labeled nucleotides as previously described (42) (Fig. 2A). Transcriptional activity increased roughly linearly over a ∼7fold range of initial concentrations. This linear increase in the overall rate of transcription dropped off at higher concentrations of components (Fig. 2A, B). High levels of TFAM led to quenching of the transcription reaction, while lower concentrations of POLRMT relative to TFAM/TFB2M reduced RNA production (Fig. S2). By using 1:1:1 stoichiometries of proteins at approximately 10X the molar concentration of template DNA, we were able to produce significant amounts of RNA (Fig. 2, Fig. S2). These conditions differ from physiological conditions, in that estimates suggest a molar ratio of TFAM:mtDNA of ∼1,000:1 *in vivo* (27), and similarly, TFAM is present in excess of TFB2M and POLRMT (43). Moreover, we use a short 0.5 kb linear DNA template as longer templates require additional protein factors (31).

**Figure 2:**
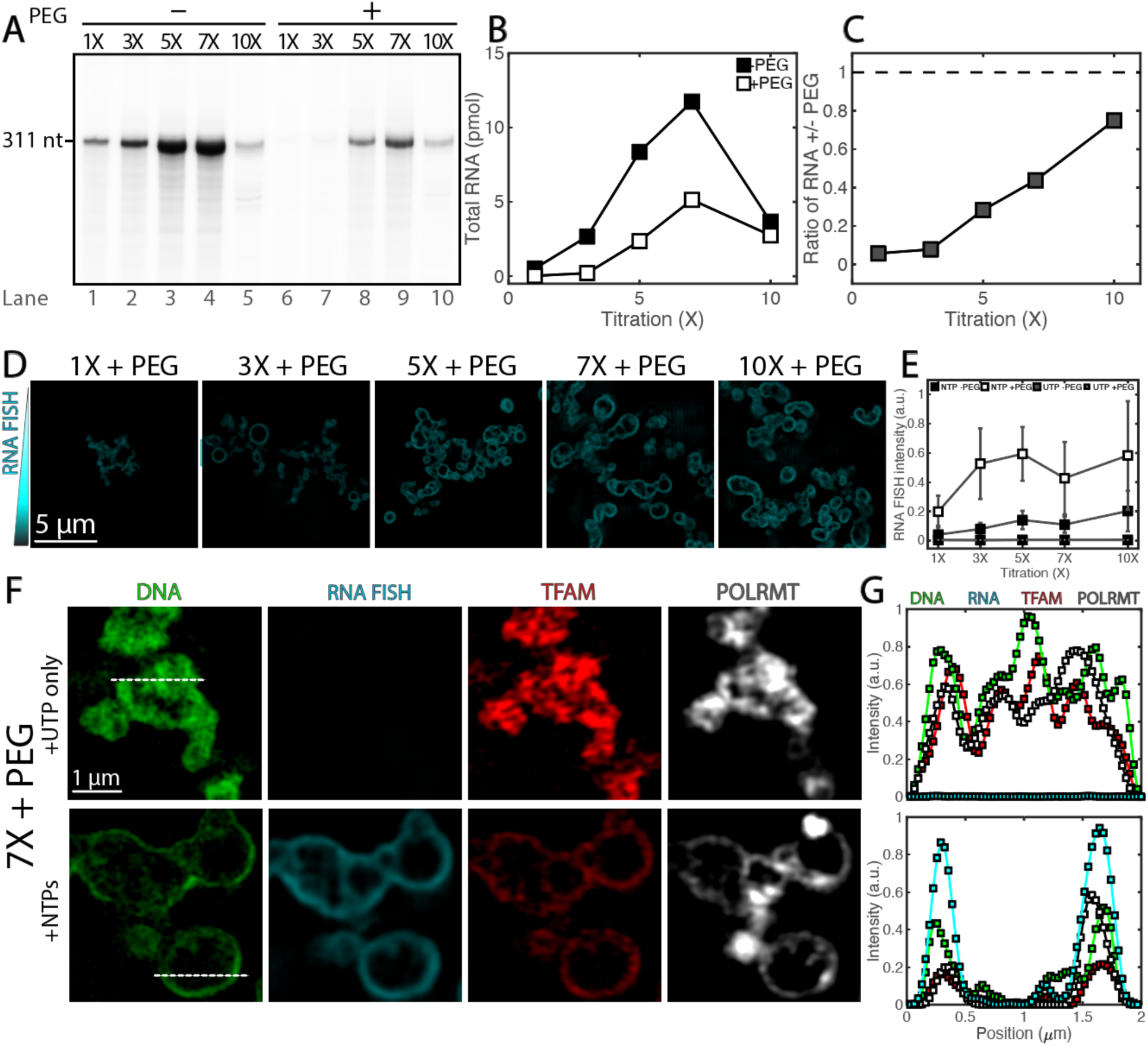
*In vitro* transcription under soluble and condensed conditions leads to changes in condensate organization. (A) RNA production rates at soluble and condensed states. The transcription run-off assay was performed using the increasing concentrations (1X to 10X) of the transcription initiation complex in the absence (lanes 1-5) or presence (lanes 6-10) of PEG. Note that the apparent decrease of the transcription efficiency in lanes 5 and 10 was due, in part, to the dilution of [α-^32^P] UTP with “cold” UTP (see SI Appendix). All reactions had equal amounts of unlabeled NTPs. (B) Quantification of RNA product from (A) as a function of reactant titration where 0% PEG is filled squares and 5% PEG is open squares. (C) Ratio of RNA production of crowded (5% PEG) to soluble (0% PEG) as a function of reactant titration. Values in A-C illustrate a representative experiment. (D) SIM images of reactions after fixation and RNA FISH, following 1 hour of reactions under the same conditions as (A). Scale bar = 5 µm. All images are at the same contrast settings. (E) Quantification of RNA FISH intensity under soluble (0% PEG) or condensed conditions (5% PEG) for the reactions (NTPs) and of the negative control (UTP). n = 3 experimental replicates, values represent averages and error bars represent standard deviation. (F) SIM images of core transcription components in condensates at 7X and 5% PEG for negative control (UTP only) and for reactions (NTPs), where DNA is in green, RNA FISH is in cyan, TFAM is in red, and POLRMT is in gray scale. Scale bar = 1 µm. Dashed lines indicate line profile. (G) Smoothed line profile for all components from (F).

Based on the optimization of mt-transcription under soluble conditions, we sought to reconstitute mitochondrial transcription under condensate forming conditions by proportionally increasing protein and DNA concentrations and including a crowder (+PEG). The condensates that form in the presence of the four different macromolecules (TFAM, TFB2M, POLRMT, and DNA) in the presence of PEG were transcriptionally active in the presence of nucleotides (Fig. 2A). Interestingly, we found the transcriptional output of condensates was 1.3 – 20-fold lower than the corresponding outputs when condensates did not form (Fig. 2B, C). Decreased rates of transcriptional output in condensates occurred most significantly first at 1X concentrations of components and approached unity with increasing reactant concentration (Fig. 2A-C).

To confirm the formation of condensates under conditions that support transcription and to relate condensate structure to function, we compared the morphologies of condensates under different conditions (Fig. 2D, E; Fig. S3). In the absence of crowder, which corresponds to the most dilute case (1X, –PEG), we did not observe condensates (Fig. S3B-E). Under transcriptionally competent, condensate-forming conditions (1X-10X, +PEG), RNA and all transcriptional components localized to the periphery of condensates, forming vesicle-like morphologies as visualized after 60 minutes of reaction (Fig. 2D; Fig. S3B-E, ref. (44)). In these vesicles, DNA tended to associate with the outermost shells, whereas RNA and proteins co-localized in the inner shell (Fig. S3F, H). The peripheral localization of the transcription machinery appears to be a consequence of active transcription, as identical, but transcription incompetent, condensates generated in the presence of only UTP nucleotides tended to retain their filled, non-vesicular, droplet-like structures (Fig. 2E-G; Fig. S3G, H; see below). These results demonstrate transcriptional activity in reconstituted mitochondrial condensates and that transcription is dampened in condensates compared to in solution.

### Newly synthesized RNA transcripts shape condensate structures

Newly synthesized RNA transcripts localize to the periphery of mt-transcription condensates (Fig. 2D). To determine whether nascent RNA is exclusively produced at the edge of the condensates or is generated internally and accumulates over time at the periphery, we performed time-course experiments (Fig. 3). RNA can be detected as early as 5 minutes in the condensate interior (Fig. 3A, B). At early time points of 5 and 10 minutes all components of the transcriptional machinery localize throughout the interior of the condensate (Fig. 3A, B). In contrast, at intervals of 20, 40, and 60 minutes, we detected a significant change in the organization of the condensate, whereby pronounced vacuoles start to appear within the condensates, coinciding with the formation of a peripheral ring containing the RNA and transcription components and increased vacuole size with time (Fig. 3A, B). All the components become peripherally located with increasing reaction time, and these morphological changes were concomitant with production of RNA (Fig. 3A, B).

**Figure 3:**
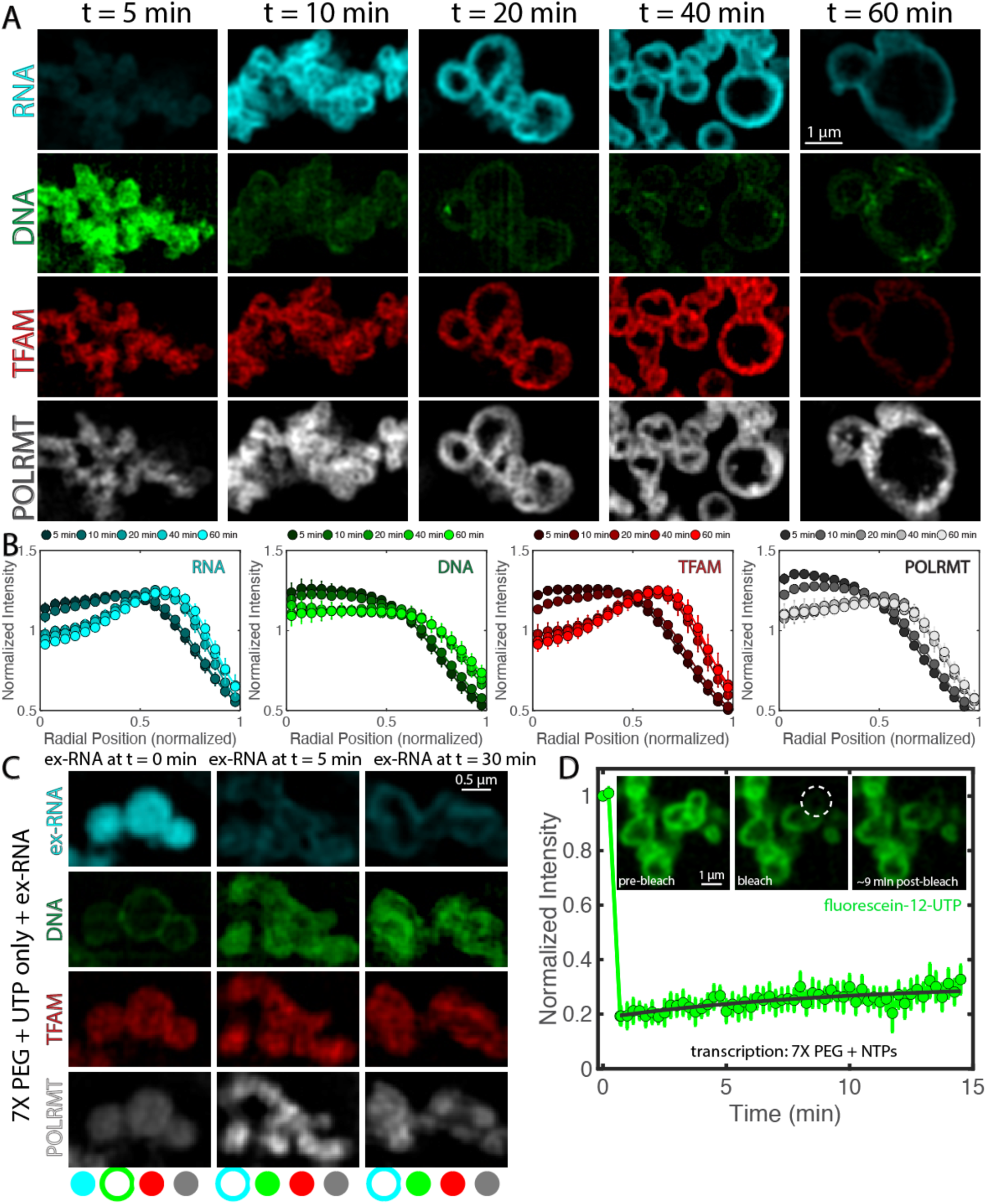
Dynamics of transcriptional condensate organization. (A) Time course of condensate morphology under reaction conditions (7X PEG, 5% PEG). Condensates were fixed and image after t = 5, 10, 20, 40, or 60 minutes at 35ºC. Single channels for RNA (cyan), DNA (green), TFAM (red), and POLRMT (grayscale) are shown. n = 4 experimental replicates and scale bar = 1 µm. Intensity of green channel was adjusted for visibility due to decrease in DAPI signal with time; all other channels are set at the same contrast settings.(B) Quantification of intensity profile of each component in the condensate, where r = 0 is the center of the condensate and r = 1 is the normalized perimeter, for each channel in (A). Shading indicates the time point corresponding to the average line profile, where darker colors are early time points and lighter colors are late time points. n = 4 experimental replicates and error bars = standard error of the mean. (C) Organization of condensates after addition of exogenous RNA (see SI Appendix) to non-reacting droplets at t = 0 (RNA added before all other proteins/DNA), t = 5 minutes (RNA added after condensates assembled 5 min at 35ºC) or t = 30 min (RNA added after condensates assembled 30 min at 35ºC). Buffer was the same as that used in the negative control (8 mM UTP). Condensates were fixed onto coverslips after 1 hour of incubation at 35ºC. n = 3 experimental replicates and scale bar = 0.5 µm. (D) FRAP recovery for transcribing droplets (NTPs, each 2mM) for 7X and 5% PEG conditions. Inset shows condensates pre-bleach, bleach, and 9 min post-bleach. Dashed circle represents region that was bleached. Scale bar = 1 µm. n = 9 droplets and error bars = standard error of the mean.

The observed reorganization of transcription components during the reaction suggests that the presence of RNA, and its increase in concentration over time, led to structural changes of the condensates. To further test the role of the newly synthesized RNA in determining condensate structure, we added exogenous RNA (ex-RNA) comparable in sequence and length to the mitochondrial components at various time points in the presence of only UTP. The goal was to mimic interactions that arise from the presence of RNA in the bulk despite the absence of transcriptional activity. Addition of ex-RNA at the beginning of mixing resulted in condensates with a layered structure, where ex-RNA, TFAM and POLRMT were in the interior, surrounded by a shell of DNA (Fig. 3C). In contrast, addition of ex-RNA after the DNA and protein components had mixed and assembled into condensates (t = 5, 30 min) resulted in condensates that had the reverse layered organization: DNA, TFAM, and POLRMT were in the interior, surrounded by a peripheral shell of ex-RNA (Fig. 3C). Here, peripheral localization of ex-RNA suggested ex-RNA coated pre-formed, existing protein-DNA-rich condensates.

We found that across these conditions, TFAM and POLRMT tended to partition with ex-RNA, suggesting that these protein-RNA interactions are energetically favorable. However, in all these cases, there was little colocalization of DNA with RNA, implying that DNA and RNA repel each other and that DNA-RNA interactions are energetically unfavorable. Indeed, combining only DNA and RNA yielded condensates with layered organization, where DNA was internal and surrounded by a shell of RNA, suggesting inherent immiscibility between DNA and RNA (Fig. S4E-F). The observation that RNA is prone to forming irregular structures on its own supports that proteins mediate the miscibility between DNA and RNA in transcriptional condensates (Fig. S4E-F).

These observations, showing that localization of components within condensates depend on the order in which different components are added, suggest that active transcription contributes to shaping condensate structure. The distinct morphologies of condensates depending on when RNA is added further indicate that active mt-transcriptional vesicular condensates represent nonequilibrium, dynamically arrested structures.

Measurements by fluorescence recovery after photobleaching (FRAP) were used to measure the mobility of actively transcribed RNA molecules by the detection of fluorescently labeled nucleotides (fluorescein-12-UTP). After ∼30 min of reaction, minimal recovery occurred over the course of ∼15 minutes (Fig. 3D). As a control, free nucleotides in a transcription-incompetent condensate rapidly exchanged resulting in only limited bleach depth (Fig. S4). Taken together, the slow internal dynamics of newly synthesized RNA molecules support our conclusion that vesicular structures formed by *in vitro* reconstitutions are dynamically arrested condensates.

### Organization of mt-nucleoids is altered upon depletion of core transcription components *in vivo*

In intact cells, mt-nucleoids typically assemble into membraneless, nucleoprotein complexes of ∼100 nm in diameter that are distributed throughout the mitochondrial network (Fig. 4A) (27). The mt-nucleoids are sites of transcription, and the newly transcribed polycistronic mtRNA localizes to adjacent mitochondrial RNA granules for further processing (10, 45). To probe structurefunction relationships of mitochondrial condensates *in vivo*, we depleted key mitochondrial transcription components and assessed their effects on mt-nucleoid fine-structure.

**Figure 4:**
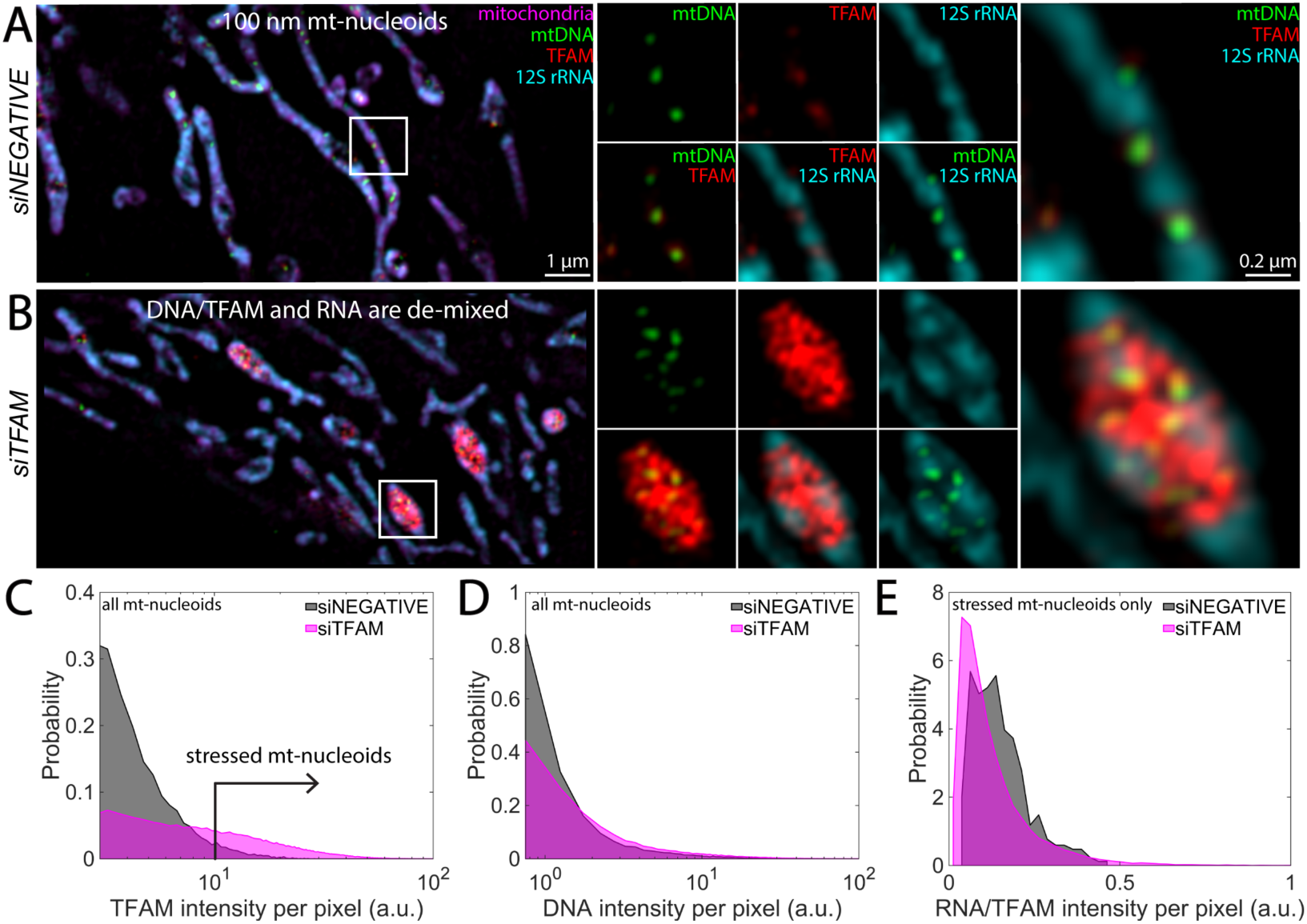
Depletion of mt-transcription components *in vivo*. (A,B) SIM images of mitochondrial components after 72 hours of siRNA treatment: siNEGATIVE (A) and siTFAM (B). Leftmost panels are the merged image of the zoomed-out version of the mitochondrial network, where mitochondria are in magenta (MitoTracker Red), mtDNA is in green (anti-DNA), TFAM is in red (anti-TFAM), and 12S mt-rRNA is in cyan (RNA FISH). Scale bar = 1 µm. White box indicates region of interest (ROI). Middle panels are signal or two-channel overlays of the ROI. Rightmost panels are the three-channel overlay for the ROI. Scale bar = 0.2 µm. Intensities across each channel are matched across siRNA conditions. n = 4 independent experimental replicates. (C) Probability distribution of TFAM pixel intensity within a segmented nucleoid, where gray = siNEGATIVE and magenta = siTFAM. Nucleoids containing higher than normal TFAM intensities are considered to be stressed as indicated by black arrow. (D) Probability distribution of DNA pixel intensity within a segmented nucleoid, where gray = siNEGATIVE and magenta = siTFAM. (E) RNA/TFAM intensity for stressed nucleoids (see arrow in D). Data represent n= 4 independent experimental replicates that were pooled together.

While complete knockdown of TFAM is embryonically lethal, partial knockdown of TFAM by RNAi (Fig. S4; Fig. S5; Fig. S6) leads to a significant reorganization of mt-nucleoids as previously observed in TFAM heterozygous knockout mice and analogous RNAi experiments (46, 47). The number of mt-nucleoids per cell, based on staining for TFAM, was reduced significantly (Fig. S5). In line with prior observations of induction of a stress response involving increased interferon stimulated gene expression and enhanced type I interferon responses upon loss of TFAM (46), dramatically enlarged clusters of mt-nucleoids were observed in TFAM depleted HeLa cells (Fig. 4B). Interestingly, these remaining stress-induced mt-nucleoids resembled the heterogeneous condensates observed *in vitro* (9), where purified TFAM and mtDNA form >1 micron-sized, multiphase droplets. Indeed, these enlarged mt-nucleoids allowed us resolve the spatial organization of the mitochondrial (ribo)nucleoprotein complexes (Fig. 4B): mitochondrial RNA localized peripherally, de-mixed from TFAM and mtDNA (Fig. 4B, Fig. S6), similar to *in vitro* condensates we formed when ex-RNA was added at later time points (Fig. 3C), and supporting the conclusion that mt-nucleoids and mtRNA exist as spatially distinct phases in live mitochondria (10, 48).

We further noticed altered phase behavior upon perturbation of other mt-nucleoid components involved in transcription (Fig. S5). The mt-nucleoid-associated protein MTERF2 (mitochondrial transcription termination factor 2), is an abundant mt-nucleoid protein, present at ∼100:1 copies relative to mtDNA *in vivo* (49). Partial knockdown of MTERF2 also led to a stress response with significantly reduced cell number and altered mt-nucleoid number (Fig. S5), associated with a pronounced population of swollen mitochondria. These mitochondria corresponded with ring-like accumulation of RNA puncta alongside the rounded membrane (Fig. S6A), reminiscent of the RNA peripheral localization of active *in vitro* transcription condensates (Fig. 2; Fig. 3). TFAM accumulated in these mitochondria and appeared to wet the inner surface of 12S RNA foci. However, mtDNA remained organized as ∼100 nm puncta surrounded by TFAM that were frequently positioned adjacent to bright puncta of 12S RNA (Fig. S6), which further supports the idea of coexisting (ribo)nucleoprotein phases. In contrast, depletion of mtDNA achieved using a mitochondrially targeted endonuclease (50), led to complete dissolution of mt-nucleoids, including TFAM, and reduction of 12S RNA signal (Fig. S6). The dissolution of DNA- and RNA-rich condensates in the mitochondrial matrix upon mtDNA depletion suggests that mtDNA nucleates mitochondrial transcriptional condensates in live cells.

Taken together, our results support the notion that the (ribo)nucleoprotein complexes in the mitochondrial matrix represent coexisting phases, whose organization is modulated by the physicochemical nature of the constituents as well as functional activity of the mt-nucleoid, specifically RNA production during active transcription.

### Computational simulation of transcription-mediated reorganization

To synthesize all our findings into a model that describes the collective phase behavior of active transcriptional condensates, we used computational modeling to recapitulate the effect of RNA production on phase behavior of mt-nucleoids. Using the simulation engine LaSSI (51), we performed Monte-Carlo (MC) simulations of coarse-grained (CG) models of binary and higherorder mixtures to probe the effects of RNA production on equilibrium condensates (Fig. 5). To preserve the overall length-scales and interaction hierarchies of the macromolecules involved in transcriptionally active condensates in the simulations, DNA molecules were modeled as chains of twenty beads, RNA as chains of ten beads, TFAM and TFB2M as chains of four beads, POLRMT as chains of three beads, and crowders as chains of four beads (Fig. 5A).

**Figure 5:**
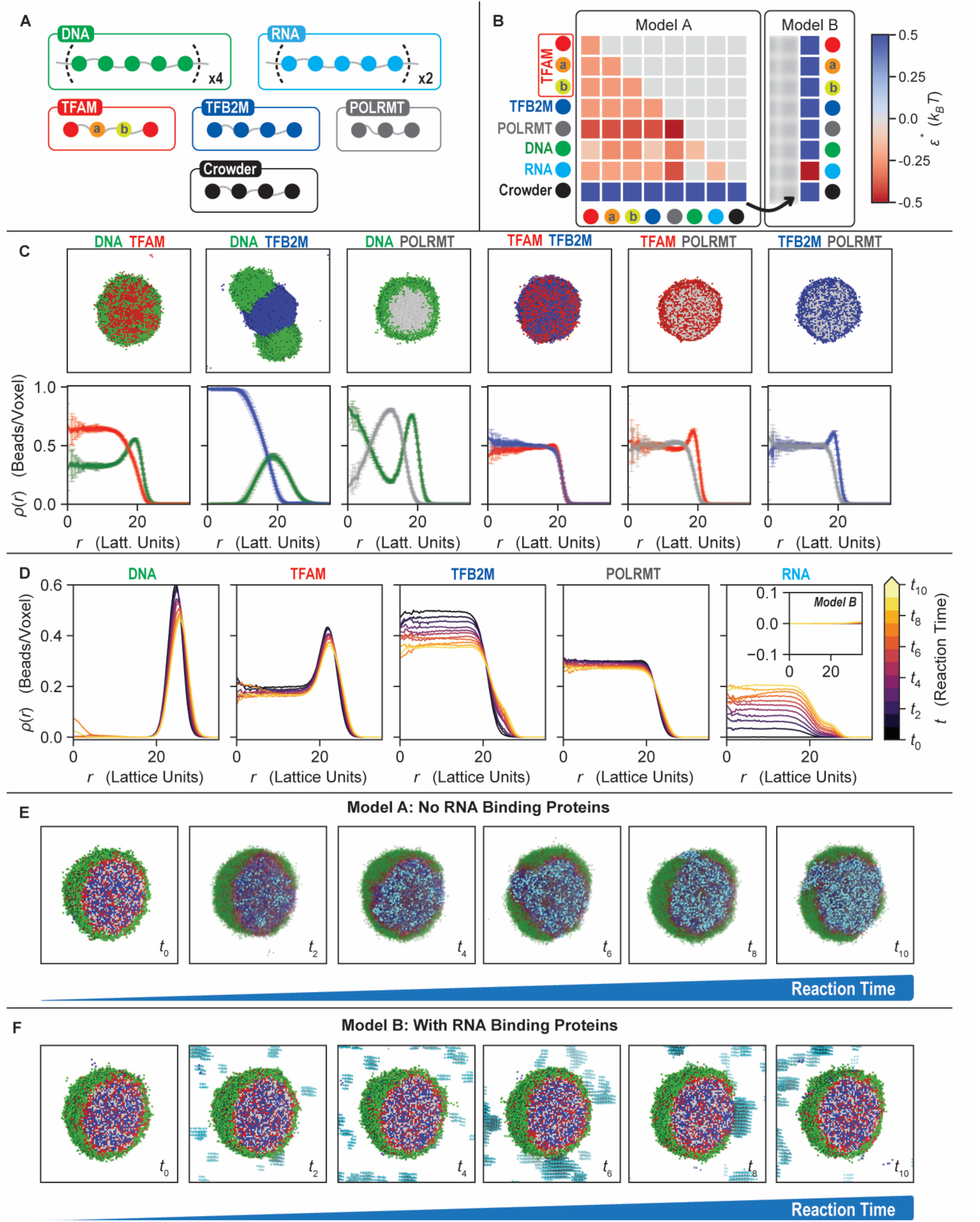
Computational modelling of mitochondrial transcriptional condensates. (A) Coarse-grained model for the mitochondrial transcriptional components. All beads are connected by implicit linkers of 2-lattice sites. DNA is modeled as chains of 20 beads. Since the mtDNA template used in the experiments produces RNA about half as long as the DNA, the RNA is a chain of 10 beads. TFAM is modeled as 4 beads (‘*X-a-b-X’*) where the central two beads, denoted as beads *a* and *b*, interact more favorably with DNA. To account for TFAM’s weak dimerization upon DNA binding, bead *b* has an additional interaction energy (–2 k_B_T) for another *b* bead, yielding a local anisotropic interaction. TFB2M is represented as 4 beads, and POLRMT is represented as 3 beads due to the highly folded nature of the molecule. Lastly, the Crowder is represented as a chain of 4 beads. (B) Interaction matrices for the two models considered. Model A lacks RNA-binding proteins, and recaptures the organization of the condensates seen *in vitro*. The Crowder has repulsive interactions with every species, including itself. The DNA and RNA have no interactions, while the rest of the molecules have favorable interactions. Model B mimics the effective inclusion of RNA-binding proteins by making the RNA-Crowder interactions to be favorable, modeling an effective RNA binding protein. (C) Representative snapshots and density profiles for the binary mixtures shown in Fig. 1C. For clarity, the Crowder is not shown. Model A qualitatively captures the experimental images. DNA & TFAM form mixed condensates with TFAM having smaller droplets within. DNA & TFB2M generate droplets that do not fully colocalize but stay closely associated. TFAM & TFB2M form well mixed droplets. POLRMT mixtures have dense POLRMT regions covered by the other component. (D) Density profiles of each component with different RNA amounts. DNA is pushed outwards as RNA is increased. For TFAM the interior most density does not change but the density at the outer rim shared with DNA is reduced. TFB2M is significantly pushed outwards as RNA is increased. RNA is accumulated evenly inside the interior of the droplet as RNA is increased. The inset corresponds to Model B and shows that no RNA is accumulated inside the condensate when simulated RNA binding proteins are present. (E) Representative snapshots of the condensates with increasing RNA for Model A. A higher amount of RNA in the system corresponds to a later time in the transcriptional reactions. For clarity, the Crowder is not shown, and non-RNA components are made transparent. The RNA evenly distributes itself inside the condensates and continues to be accumulated as the reaction continues. (F) Representative snapshots like (E) but for Model B. With a suitable RNA binding protein, the RNA can be prevented from going inside the condensates.

We first parameterized the contact energies between pairs of molecules by reproducing experimentally observed morphologies of single and binary mixtures (Fig. 5C, compare to Fig. 1B, C) (9). The experimentally measured colocalization and condensate homogeneity were used as proxies for stronger heterotypic interactions, while spatial inhomogeneities within the condensates were used as proxies for stronger homotypic interactions. TFAM has an additional anisotropic interaction to account for its weak dimerization upon DNA binding (Fig. 5B) (52). Simulations of the binary mixtures of transcriptional condensate components generated morphologies (Fig. 5C; Fig. S7A) with spatial organizations, quantified in terms of radial density profiles, that recapitulate the experimental results (Fig. 1C). For example, TFAM-DNA and POLRMT-DNA formed multi-phase droplets in the simulations, while TFB2M and DNA behaved as distinct coexisting phases, and pairs of proteins tended to form well-mixed droplets, as observed *in vitro* (Fig. 1C, Fig 5C).

Extending this modeling approach to the quaternary condensates, we find the formation of heterogenous droplets that are consistent with our experimental results (Fig. 5E, t=0; compare to Fig. 1D). Simulations show the formation of layered droplets with DNA being localized almost exclusively at the periphery (Fig. 5D; Fig. S7B). The three proteins tend to be enriched in the interior of the droplet, enveloped by a shell of DNA. This multi-phase organization is consistent with the heterogeneity and co-localization of components in the four-component dynamically arrested droplets observed *in vitro* (Fig. 1D).

Based on our experimental observations of reorganization of transcriptional components upon synthesis of nascent RNA in the condensate, we sought to explore how transcription affects the morphologies of condensates *in silico*. At the initial time point *t*_0_, all reactants are fully mixed, and no RNA is present, while at some later time, *t*_*i*_ > 0, the system contains RNA, [*R*]_*i*>0_; conversely, if the system has an amount of RNA, [*R*]_*i*_, then the system is at *t*_*i*_. Therefore, titrating the amount of RNA in the simulations allowed us to model the equilibrium structures of condensates that should form when transcription occurs (Fig. 5). We find that owing to the favorable interactions between the RNA and the proteins, newly synthesized RNA is readily incorporated into the existing droplet (Fig. 5E). RNA distributes itself into the interior of the droplets (Fig. 5D; Fig. S7B), and with increasing levels of RNA, TFB2M is peripheralized from the interior, which further pushes DNA to be localized almost exclusively to the periphery (Fig. 5D). Therefore, in accord with experimental observations, the simulations show that transcription directly remodels the organization of the condensate components. Strikingly, the computationally predicted equilibrium structures are exact facsimiles of structures we obtain upon mixing ex-RNA with components early on, t=0 (Fig. 3C).

While we find that RNA accumulates in vesicular condensates upon transcription *in vitro*, this organization is not observed *in vivo*. It is likely that in non-stressed cells newly synthesized RNA molecules are bound by RNA-binding proteins and processed in mitochondrial RNA granules, which are phase separated structures that are often located adjacent to mt-nucleoids (10, 45). To test whether the action of RNA-binding proteins may account for the distinct RNA distribution in DNA-rich condensates *in vitro* and *in vivo*, we incorporated an additional favorable interaction between the crowder and the RNA in our simulations to mimic the effects of association of RNAbinding proteins to the newly synthesized RNA under steady-state conditions *in vivo* (Fig. 5B). Under these conditions, the reorganization of transcriptional components mirrors their distribution *in vivo* (Fig. 4; Fig. S5; Fig. S6), where RNA no longer accumulates within the DNA-rich droplet but condenses separately in the bulk (inset Fig. 5D, E; Fig. S7C). Overall, these simulations recapitulate the *in vivo* behavior of mitochondrial condensates, and they also affirm the inference that vesicular structures observed *in vitro* are non-equilibrium, dynamically arrested phases.

## Discussion

We report here the reconstitution of transcriptionally active, multi-phasic condensates using the human mitochondrial transcription machinery as a model system. We show that, when compared to bulk reactions realized in the absence of condensates, the transcriptional rate is reduced under condensate-forming conditions. This, we attribute to the dynamically arrested nature of transcriptionally active condensates that form *in vitro*. We also observe that the production of RNA alters the spatial organization of condensates, thus providing direct evidence for a dynamic interplay between the structure of condensates and the functional activities they harbor.

Decreased rates of transcription within condensates that form *in vitro* are likely associated with slower internal dynamics that we observe in these condensates. These may be thought of as increasing the Damköhler numbers (*D*_*a*_ *∼ k/D*), which compare reaction rates (*k*) relative to the diffusive mass transfer rates (diffusion coefficient, *D*) (53). Whereas in bulk solution, reactants can diffuse quickly, and the rate of transcription is only limited by the speed of the polymerase (*Da* << *1*, reaction-limited), the significantly slow dynamics associated with the condensate environment, as they exist in mitochondrial condensates *in vivo*, suggest that condensates experience diffusion- or transport-limited kinetics (*Da >> 1*, diffusion-limited) (11). Our rough estimates suggest that less than one round of transcription occurs under condensate-forming conditions indicating that not all DNA templates were actively transcribed. This behavior is in line with the situation in live cells, where only a minority of <5% of the mitochondrial nucleoids are actively transcribed at any given time (54, 55). Comparisons of *in vitro* and *in vivo* mitochondrial transcription properties and phase behavior are complicated by difference in the stoichiometry and DNA templates required to reconstitute efficient mt-transcription *in vitro* under both soluble and condensate conditions. For example, *in vivo* TFAM is present in roughly 1,000 copies per copy of mtDNA (27) and in significant stoichiometric excess of TFB2M and POLRMT (43), whereas *in vitro* reconstitution systems require use of roughly equimolar ratios of proteins to generate detectable RNA product as relatively high TFAM levels significantly quench the reaction (Fig. S2) (30). Moreover, the mitochondrial genome is circular and 16 kb in size, whereas *in vitro* transcription assays use short linear DNA templates of ∼0.5 kb, since longer transcripts cannot be efficiently generated *in vitro* without the presence of other protein factors (31). The RNA molecules generated *in vivo* are long transcripts that are quickly bound and modified by RNAbinding proteins (56), whereas our *in vitro* system generates short (∼300 nt) RNAs in the absence of any such RNA-modifying proteins.

There are growing reports of transcription occurring within condensates *in vivo*. RNA Pol I transcribes rRNA in the multiphase nucleolus (7, 57), RNA Pol II has been reported to produce mRNA in transcriptional condensates (58, 59), and POLRMT generates long, polycistronic mtRNA in mitochondrial condensates (9, 10, 31). While condensate formation is not an absolute requirement for transcription (60), there is growing evidence that condensate formation may offer unique advantages for regulation of transcription *in vivo*. First, the condensed phase enriches for specific reactants, which may regulate mass action effects (11). For example, SUMOylation can be increased ∼36 fold in engineered condensates *in vitro* (61). In addition, increased dwell times of proteins associated with the condensate microenvironment may also be conducive for assembly of reacting complexes (62). In support of this hypothesis, an early FRAP study showed that the kinetic behavior of multiple RNA Pol I components could only be explained by inclusion of a slow kinetic component prior to binding of the polymerase subunits to the promoter, possibly reflecting slowed diffusion in a nucleolar condensate (63).

Importantly, we find that the structure of the transcriptional condensate is affected by its activity. Production of a new chemical species–RNA–in the otherwise DNA- and protein-rich transcription condensates leads to non-equilibrium changes in condensate organization, reflected by the emergence of vesicle-like morphologies. Intriguingly, similar vesicle morphologies have been observed for simple *in vitro* RNA-protein systems (44), *in vivo* liquid spherical shells of the DNA- and RNA-binding protein TDP-43 (64), and *in vitro* liquid spherical shells of DNA and poly-LLysine (65). RNA has also been shown to form a corona on the surface of engineered condensates, directly impeding their coarsening (66). Vesicular structures are likely to be thermodynamically accessible in specific regions of protein-nucleic acid phase diagrams. Importantly, while equilibrium simulations predict the organization of multiphase condensates at the onset of the transcription reaction, they do not produce vesicle-like structures as transcription progresses. This reinforces our conclusion that the vesicles we observe *in vitro* are non-equilibrium structures.

The vesicle formation observed due to RNA generation in transcriptionally active condensates *in vitro* differs from that of the canonical mt-nucleoid organization in mitochondria *in vivo* (27, 28), where mt-nucleoids remain as condensed, ∼100 nm droplet-like structures, and mtRNA localizes to separate mtRNA-processing granules that coat the mitochondrial inner membrane and are demixed from the mt-nucleoid (10). Our simulations, which are based on experimentally determined interaction parameters, suggest that during transcription, mtRNA is tethered to the mt-nucleoid as it is being transcribed by POLRMT but later moves – be it actively or passively – out of the mt-nucleoid into more energetically favorable mtRNA-processing granules in mitochondria. An attractive model, consistent with our observations, is that *in vivo* the nascent mtRNA is effectively handed off from the mt-nucleoids condensates, from where it is generated, to the more energetically favorable mitochondrial mtRNA-processing granules, which are themselves condensates (10). This scenario is supported by our computational simulations which demonstrate that the presence of RNA-binding activities outside the condensate is sufficient to remove the accumulating RNA from the condensate.

Finally, our observations regarding mt-transcription condensates are also relevant to other cellular transcriptional processes. Transcription of ribosomal DNA (rDNA) by RNA Pol I occurs in fibrillar centers (FC) in the nucleolus, which unlike the mt-nucleoid, contains enveloping coexisting phases of RNA and proteins (7). However, both mt-nucleoids and FCs represent cores of a DNA-rich phase (mtDNA or rDNA, respectively), in which the nascently transcribed RNA moves radially outwards, into adjacent mtRNA granules in mitochondria (45) or towards the dense fibrillar component (DFC) and granular components (GC) in the nucleolus (57), respectively. Similarly, RNA Pol II can self-assemble with the transcriptional machinery to form condensates, particularly at super-enhancers, in the mammalian nucleus (58). Interestingly, for RNA Pol II condensates, rapid, local RNA production has been shown to result in complete dissolution of the condensate, underscoring a feedback mechanism between phase behavior and RNA production as also observed here for mitochondrial transcription (25). Similar condensation is also seen in the bacterial nucleoid, where RNAP clusters with transcription factors at specific sites, particularly rDNA, in the bacterial genome (67, 68). The common observation that emerges from these observations is that RNA is not retained in DNA-rich phases, pointing to an intrinsic energetic barrier for their mixing.

## Materials and Methods

Description of protein purification, phase separation assays, *in vitro* transcription reactions, cell culture and knockdown experiments, and computational modelling is detailed in full in the SI Appendix.

## Acknowledgments

We are grateful for discussions with Misteli group members, J. Jones and M. Taylor for protein expression and purification of TFAM at the NIH/NCI/CCR Protein Production Core; and T. Karpova and D. Ball for assistance with Structured Illumination Microscopy and Laser Scanning Confocal Microscopy as part of the NIH/NCI/CCR Optical Imaging Core. FD and RVP are also grateful to K. M. Ruff for many helpful and clarifying discussions. Research in the Misteli lab is supported by funding from the Intramural Research Program of the National Institutes of Health (NIH), National Cancer Institute (NCI), and the Center for Cancer Research (CCR) (1-ZIABC010309); MF was supported by a Postdoctoral Research Associate Training (PRAT) fellowship from the National Institute of General Medical Sciences (NIGMS, 1Fi2GM128585-01). DT is supported by NIH R35 GM131832. This work in the Pappu lab was supported by grants from the National Institutes of Health (R01NS121114, 5R01NS056114), the Air Force Office of Scientific Research (FA9550-20-1-0241), and the St. Jude Collaborative Research Consortium on the Biology of Membraneless Organelles.

## Data and materials availability

All data and materials are available from authors upon reasonable request.

## Supplementary Materials

Materials and Methods

Figures S1-S7

## Notes

**Competing Interest Statement:** MF, FD AS, DT and TM declare no conflict of interest. RVP is a member of the Scientific Advisory Board of Dewpoint Therapeutics Inc. The work here was not supported or influenced by this affiliation.

### Competing Interest Statement

MF, FD AS, DT and TM declare no conflict of interest. RVP is a member of the Scientific Advisory Board of Dewpoint Therapeutics Inc. The work here was not supported or influenced by this affiliation.

### Summary of Updates

We have added more supplementary data, and we have updated the text accordingly.

