## Supplementary Information for "Structure-function relationships in mitochondrial transcriptional condensates"

**This PDF file includes:**

Supplementary Materials and Methods
Fig. S1 to S7
Supplementary References

### Supplementary Material and Methods

#### Cell culture

HeLa cells (ATCC, CCL-2, Lot. #70000153) were grown at 37C and with 5% CO2 in Dulbecco's Modified Eagle Medium (DMEM, Thermo Fisher Scientific, #11960-044) supplemented with 10% Fetal Bovine Serum (Thermo Fisher Scientific, #10437), 1% glutamine (Thermo Fisher Scientific, #25030081), and 1% penicillin/streptomycin (Thermo Fisher Scientific, #15140-122).

#### siRNA

Silencer Select Pre-Designed siRNA constructs were used to knock-down mitochondrial proteins (TFAM–s14002, POLRMT–s10826, TFB2M–s34561, TEFM–s36219, mTERF1–s15508, mTERF2–s37162, mTERF3–s27208, mTERF4–s43559). Cells were seeded on high precision cover slips (Azer Scientific) in 12-well plates ($2.5 \times 10^4$ cells) or in 96-well plates ($2.5 \times 10^3$ cells) on Day 0. On Day 1, cells were transfected with siRNA (15 nM final) using DharmaFECT transfection reagent (Horizon Discovery) and incubated for 72 hours. As a negative control, Silencer Select Negative Control No. 2 siRNA was used (Thermo Fisher Scientific, ASO2GNFA). As a positive control of siRNA transfection, AllStars Hs Cell Death siRNA (Qiagen, 1027299) was used.

#### Western Blot

Trypsinized cells were collected from 12-well plates, centrifuged, washed with ice-cold PBS, dissolved in 2X Laemmli Sample Buffer (Bio-Rad), and denatured for ~10 min at 95ºC. Samples were stored at -20ºC. Protein concentration was estimated using a Bradford assay. Equal protein amounts of each sample were loaded onto a 10-well NuPAGE 4-12% Bis-Tris Protein Gel at 150 V for 1 hour followed by wet transfer onto a membrane at 200 mA for 2 hours. Membranes were blocked with 5% BSA and 1X TBST for ~30 minutes at room temperature and incubated with primary antibodies (TFAM (Sigma, HPA063684), beta-actin (Sigma, A2228), MTERF2 (Thermo Fisher Scientific, PA5-109989), and tubulin-alpha (Bio-Rad, VMA00051)) in 5% BSA and 1X TBST at 4ºC overnight followed by three wash steps. Membranes were incubated with HRP secondary antibodies for 1 hour at room temperature followed by three wash steps. Proteins were detected using the Amersham ECL Western Blotting Detection Reagent on a Bio-Rad ChemiDoc Imager and quantified with ImageJ.

#### mtDNA depletion

Cells were seeded on Day 0 on coverslips on high precision cover slips (Azer Scientific) in 12-well plates ($2.5 \times 10^4$ cells) or in 96-well plates ($2.5 \times 10^3$ cells). On Day 1, media was supplemented with 1 mM sodium pyruvate (Thermo Fisher Scientific) and 50 ug/ml uridine (Thermo Fisher Scientific) (*43*). Cells were transfected with a plasmid containing the DNase Herpes Simplex Virus UL12.5M185 (pMA4008, Addgene plasmid #70109) using the FuGene transfection reagent and incubated for 72 hours (*43*).

#### Fixation and labelling

Cells were incubated with 100 nM MitoTracker Red CMXRos (Thermo Fisher Scientific) for ~15 min at 37ºC and 5% $CO_2$. To fix cells, 16% paraformaldehyde (PFA) was diluted 1:2 in PBS to make an 8% solution, which was then diluted 1:2 directly into cell culture media (4% PFA final), and cells were fixed for 20 minutes at room temperature. Cells were washed with PBS and stored at 4ºC. To permeabilize cells, cells were treated with 0.1% Triton-X for 10 min and washed with PBS. Coverslips were incubated with primary antibodies (anti-DNA clone AC-30-10, EMD Millipore, #CBL186, and anti-TFAM, Atlas Antibodies, HPA063684) without blocking agents for 1 hour (1:500 dilution) at room temperature in a humidity chamber. Coverslips were then washed three times with PBS. Secondary antibodies were applied at 1:1000 for anti-rabbit-488 (Thermo Fisher Scientific, A11008) or 1:50 for anti-mouse-405 (Thermo Fisher Scientific, A31553) for 1 hour at room temperature in a humidity chamber. Cells were washed three times with PBS. To maintain adherence of antibodies, cells were fixed a second time with 4% PFA in PBS for 10 min and washed with PBS. Custom RNA FISH probes were designed to complement the 12S mt-rRNA as well as COI mt-mRNA transcript Stellaris ® RNA FISH Probe Designer (Biosearch Technologies, Inc. Petaluma, CA) available online (www.biosearchtech.com/stellarisdesigner, version 4.2) (*9*). An adapted protocol was followed based on the manufacturer's instructions available online (www.biosearchtech.com/stellarisprotocols). Cells were mounted onto glass slides with ProLong Gold Antifade Mountant (Thermo Fisher Scientific, P36930) and left to cure at room temperature for >24 hours.

**DNA template for transcription assays**
Template DNA (~500 bp) was amplified as described previously using pT7blue plasmid containing -70 to +70 of native LSP sequence (*60*) to generate ~300 bp of the transcribed downstream sequence. The reaction was carried out using PCR thermocycler with Taq 2X Master Mix (New England Biolabs) using forward U19 primer (GTTTTCCCAGTCACGACGT) and reverse RO 300 primer (CTGGAAAGCGGGCAGTG). PCR reactions were run (95ºC 2 min, 35 cycles of 95ºC 15s, 55ºC 30s, 68ºC 30s, followed by extension of 68ºC for 5 min) and were pooled together to be purified on columns from a GeneJET PCR purification kit using isopropanol precipitation to yield ~1-2 µg/µl of DNA template resuspended in 20 mM Tris-HCl, pH 7.9.

**Protein purification**
Human TFAM was purified as previously described (*9*). Briefly, mature TFAM (res 43–246) was transformed I BL21 star (DE3) pRare *E. coli.* 500 ml of bacterial culture containing Dynamite media plus kanamycin and chloramphenicol was inoculated with TFAM and incubated until OD of 7. Expression was induced with IPTG at 0.5 mM, and the culture was incubated overnight at 16ºC at 220 rpm. The culture was centrifuged, and the pellet was resuspended and lysed in lysis buffer [20 mM Tris-HCl, 500 mM NaCl, pH 8.0]. Cells were lysed with a microfluidizer and centrifuged for 30 minutes (70,000 g, 4ºC). Protein was purified on an immobilized metal affinity chromatography (IMAC) column and eluted in elution buffer [20 mM Tris-HCl, 500 mM NaCl, 250 mM Imidazole, pH 8.0], and dialyzed into lysis buffer [20 mM Tris-HCl, 500 mM NaCl, pH 8.0] and temporarily stored at 4C. Nucleic acids were removed by purifying TFAM with a HiTrap Heparin High performance column (GE Healthcare). Protein was diluted to ~ 0.2 mg/ml in Heparin buffer (20 mM Tris-HCl, 300 mM NaCl, pH 7.5). The column was pre-equilibrated with Heparin buffer, followed by loading of diluted TFAM, and washed with 5X column volumes of Heparin buffer. Protein was eluted using a two-step gradient of 650 mM NaCl followed by 1 M NaCl. Fractions enriched in purified TFAM were pooled together and 10% (vol/vol) glycerol was added prior to flash freezing with liquid nitrogen. Proteins were stored at -80ºC.
Human TFB2M (res 21-396) and human POLRMT (res 120-1230) were purified as previously described (*61*).

**Protein labelling**

Protein was fluorescently labelled using DyLight antibody labelling kits (Thermo Fisher), where TFAM was labelled with DyLight-488, TFB2M was labelled with DyLight-594, and POLRMT was labelled with DyLight-650 or -594. Fluorescent proteins were added in trace amounts to unlabeled protein (~1:100 dilutions) prior to phase separation and transcription assays.

**Equilibrium phase separation assays**

Frozen protein aliquots were thawed at room temperature and kept on ice. Proteins were concentrated and/or buffer exchanged in 40 mM Tris-HCl, pH 8.0, 5 mM BME, 500 mM NaCl using 0.5 mL Centrifugal filters 10-kDa (Amicon), as needed. Protein concentration was estimated by the Bradford Assay using BSA standards (BioRad) on a Denovix spectrophotometer. To initiate phase separation under equilibrium conditions, concentrated protein solutions at high salt (300-500 mM NaCl) and DNA (20 mM Tris-HCl, pH 7.9, 0 mM NaCl), were diluted into low-salt buffer to reach final buffer conditions of 20 mM Tris-HCl, pH 8.0, 10 mM $MgCl_2$, 20 mM BME, 100 mM NaCl and containing either 5 or 10% PEG (MW 3,350 Da, Sigma P4338). To create the final solutions, the reagents were added to an Eppendorf tube in the following order: water and/or PEG solution that also contained trace amounts of DAPI, 10X buffer (200 mM Tris-HCl, pH 8.0, 100 mM $MgCl_2$, 200 mM BME), 300-500 mM NaCl solution, protein (TFAM, POLRMT, and/or TFB2M), and/or DNA (see Table 1 or Figure Legend 1). Solutions were briefly centrifuged and pipetted up and down to ensure complete mixing. Approximately 3 ul of solution was added to Pluronic F-127 (Sigma, P2443) treated (for 1-3 component droplets) or untreated (for four component droplets) high-precision coverslips (Azer) and sealed with a 4.5 mm diameter x 0.6 mm depth silicon isolator (Grace-biolabs) and incubated for ~30-60 minutes at room temperature prior to imaging.

**Table 1 Composition of binary and ternary droplets**

| | TFAM (µM) | TFB2M (µM) | POLRMT (µM) | DNA (nM) | PEG (%) |
|---|---|---|---|---|---|
| **TFAM+DNA** | 10 | 0 | 0 | 500 | 10 |
| **TFB2M+DNA** | 0 | 10 | 0 | 500 | 10 |
| **POLRMT+DNA** | 0 | 0 | 1.5 | 500 | 5 |
| **TFAM+TFB2M** | 10 | 10 | 0 | 0 | 10 |
| **TFAM+POLRMT** | 10 | 0 | 1.5 | 0 | 5 |
| **TFB2M+POLRMT** | 0 | 10 | 1.5 | 0 | 5 |
| **TFAM+TFB2M+DNA** | 10 | 10 | 0 | 500 | 10 |
| **TFAM+POLRMT+DNA** | 10 | 0 | 1.5 | 500 | 5 |
| **TFB2M+POLRMT+DNA** | 0 | 10 | 1.5 | 500 | 5 |
| **TFAM+TFB2M+POLRMT** | 10 | 1.5 | 1.5 | 0 | 5 |

***In vitro* transcription reaction**

Reactions were set up similarly to our phase separation assays but with the incorporation of ribonucleotides (either NTPs or UTP). Briefly, water and/or PEG solution, 10X transcription buffer (200 mM Tris-HCl, pH 8.0, 100 mM $MgCl_2$, 200 mM BME), 500 mM NaCl solution, NTPs or UTP (negative control), protein (TFAM, POLRMT, and/or TFB2M), and lastly DNA. For 1X reactions, the composition was 0.6 µM TFAM, 0.6 µM TFB2M, 0.6 µM POLRMT, and 50 nM

DNA. The concentration of NaCl was as follow: ~20 mM (1X), ~40 mM (3X), ~60 mM (5X), ~80 mM (7X), and ~100 mM (10X). Final concentration of buffer in reaction conditions was: 20 mM Tris-HCl, pH 8.0, 10 mM $MgCl_2$, 20 mM BME, ~20-100 mM NaCl, 2 mM NTPs (each), 0 or 5% PEG. All other reactions were a multiple (3, 5, 7, 10X) of all reactions of the 1X reaction. Reactions were performed in a total volume of 20 ul in an Eppendorf tube on a heat block set at 35ºC for one hour. Negative control for the reactions contained the same composition, except instead of 2 mM NTPs (each), 8 mM UTP was used.

**Transcription assays using radionucleotides**

Transcription reactions were carried out using DNA template (50 nM), mtRNAP (600 nM), TFAM (600 nM) and TFB2M (600 nM) in a transcription buffer containing 20 mM Tris (pH=7.9), 10 mM NaCl, 10 mM $MgCl_2$, 20 mM $\beta$-mercaptoethanol in the presence of ATP (0.3 mM), GTP (0.3 mM), CTP (0.3 mM), UTP (0.05 mM) and 0.1 $\mu$l of [α-$^{32}$P] UTP (800 Ci/mmol). Where indicated, 5% PEG3350 (Sigma) was added to the transcription reaction. The concentration of all reaction components, except [α-$^{32}$P] UTP, was increased proportionally in 3X, 5X, 7X, and 10X reactions. To prevent protein precipitation, NaCl concentration was increased to 35 mM (3X reaction), 55 mM (5X), 80 mM (7X), and 110 mM (10X). All reactions were incubated for 30 min at 37ºC and treated with 0.2 mg/ml of proteinase K for 1 h at 55ºC. Reactions were stopped by the addition of an equal volume of 95% formamide and 0.05 M EDTA. The products were resolved by 12% PAGE containing 6 M Urea and visualized by PhosphorImager (GE Health). The amount of the transcripts produced in each reaction was quantified by ImageQuant TL software.

**Time-course for *in vitro* transcription reactions**

Reactions were set up as before but with a total larger volume of (~60 ul). After the reaction master mix was assembled, the mix was then split into ~10 ul aliquots into five separate tubes and placed on a heat block. One tube was then removed at the indicated time points: 5, 10, 20, 40 or 60 minutes and prepared for fixation and labelling (see below).

**Exogenous RNA experiments**

Synthetic RNA was transcribed using the SP6 promoter that was previously incorporated in the DNA template with a MAXIscript SP6/T7 transcription kit (Thermo Fisher Scientific). For direct labelling RNA in RNA-only droplet assays (Fig. S4), fluorescent nucleotides (CF-640R-UTP, Biotium) were added to the reaction. Synthetic RNA was purified using a RNeasy Mini Kit (Qiagen) and purified in water. Mixtures were assembled similar to the reactions as previously described, but with the addition of 8 mM UTP instead of NTPs. While setting up the mixture, RNA was added at different time points: t = 0 min (RNA added before protein or DNA), t = 5 min (RNA added after all of the components had been added and incubated on the heat block for 5 min), or t = 30 min (RNA added after all of the components had been added and incubated on the heat block for 30 min). In parallel, two controls were performed alongside these experiments. As a negative control, only 8 mM UTP was added, and as a positive control, NTPs (2 mM, each) were added to trigger the reaction.

**Droplet fixation and mt-RNA FISH labelling**

Reactions were fixed by direct addition of 16% PFA to *in vitro* transcription reactions in Eppendorf tubes, to create a final concentration of ~0.5% PFA, and the mixture was pipetted up and down several times. After ~5 minutes of incubation in the Eppendorf tube, aliquots of 3 ul were deposited to untreated high precision glass coverslips (Azer) in 12 well plates for an additional ~25 minutes to allow fixation of condensates to the glass coverslips. Reactions were washed off with PBS, leaving condensates that had become fixed to the glass coverslip and removing any soluble components in the aqueous phase. Custom RNA FISH probes (13 probes of ~20 bases) were designed to complement the mt-RNA transcript generated from the DNA template upon *in vitro* transcription using Stellaris ® RNA FISH Probe Designer (Biosearch Technologies, Inc. Petaluma, CA) available online (www.biosearchtech.com/stellarisdesigner, version 4.2). An adapted protocol was followed based on the manufacturer's instructions available online (www.biosearchtech.com/stellarisprotocols). Briefly, coverslips were washed with Wash Buffer A for ~5 minutes. Coverslips were then treated with mt-RNA FISH probes resuspended in hybridization buffer in a humidity chamber placed in a 37ºC incubator for ~2 hours. Coverslips were subsequently washed with Wash Buffer A for 30 minutes at 37ºC. Media was aspirated and replaced with DAPI in Wash Buffer A for an additional 30 minutes at 37ºC. Media was removed, and coverslips were washed in Wash Buffer B for ~5 minutes. All media was aspirated, and coverslips were sealed onto glass slides with Prolong Gold Antifade Mountant (Thermo Fisher Scientific) for >24 hours at room temperature to allow for curing before super-resolution imaging.

**FRAP experiments**

The reactions were assembled in 1.5 ml Eppendorf tubes the same as above and placed on the heat block at 35ºC. After ~30 min, the reactions were transferred to a pre-heated 8-well imaging chamber and covered with pre-heated mineral oil (Sigma, M5904) (all at 35ºC). A ~1 µm spot on the condensates was photobleached for ~30 s and images were acquired every 15 s.

**Light microscopy**

Super-resolution imaging was performed using Structured Illumination Microscopy (SIM) on untreated droplets (equilibrium conditions), fixed droplets (after *in vitro* transcription), and on fixed cells at room temperature using an ELYR PS.1 on an AxioObserver Z1 inverted microscope operating ZEN software. Samples were imaged with all four laser lines (405, 488, 561, 647 nm) and imaged with a 63X/1.4 NA oil Plan Apochromat objective. Slides containing 200 nm fluorescent beads were used to standardize for channel alignment. ZEN software was used for SIM processing and channel alignment. FRAP experiments were performed on a Carl Zeiss LSM780 microscope using a 100X objective, and droplets were imaged using transmitted light and 488/561 nm lasers. The heating control stage insert was set to 35ºC to maintain reaction conditions. For high-throughput experiments, 384-well plates containing tissue culture cells after siRNA treatment were imaged on a fully automated Yokogawa CV7000S spinning disk microscope with a 60X water immersion objective, 405, 488, 561, and 647 laser lines as maximum intensity projection images.

**Quantitative image analysis**

Images were quantitatively analyzed using MATLAB, and images were visualized using ZEN, FIJI, and/or Imaris software. To characterize morphology, droplets were segmented from background using standard Matlab Image Toolbox functions and droplet features were quantified based on size, morphology, and number. To estimate degree of miscibility between pairs of components, cross-correlation coefficients were quantified for all pixels within the droplet and were averaged across all droplets per condition.

**Bioinformatics**
Protein sequences were obtained from PUBMED, and domains were annotated using Matlab (*29*). Disorder was predicted from several algorithms using the Database of Disordered Protein Predictions (D2P2) (*62*), and graphically represented using Matlab.

**Computational modeling**
The simulations were performed using the LaSSI simulation engine (*44*). A lattice-size, $L$, of 120 was used for each simulation. The following move frequencies were used for all simulations:

**Table 2 MC move frequencies**

| Move | Frequency | *Normalized* Frequency |
|---|---|---|
| Small Cluster | 0.0081 | 1 |
| Multi-Local | 0.0163 | 2 |
| Double Pivot | 0.0813 | 10 |
| Pivot | 0.0813 | 10 |
| Chain Translation | 0.0813 | 10 |
| Snake | 0.2439 | 30 |
| Local | 0.4878 | 60 |

For more detailed information of the moves, see (*44*). There are two new moves employed here:
1. Small Cluster moves pick a random chain in the system and calculate the cluster that chain belongs to. Clustering is based on a maximal bead distance of $\sqrt{3}$. If the cluster is smaller than 2, or larger than 250, the move is rejected. Otherwise, the cluster is displaced randomly in a radius of $\frac{L}{2}$. For detailed balance, we recalculate the cluster, and if the resulting cluster is different from before, the move is rejected. Otherwise, the move is accepted.
2. Multi-Local moves pick a random bead in the system. That bead, and the beads that are covalently bonded to it are removed from the lattice. All beads are attempted to be re-placed within a 2-lattice site radius of the first bead. Steric clash or incorrect linker lengths results in move rejection. If appropriate lattice-sites are found, the standard Metropolis-Hastings criterion is used for acceptance.

***Two-component Mixtures***
For simulations of the binary mixtures, the following protocol is used. For $5 \times 10^7$ steps, the simulation temperature starts at $1000T^*$, and the following constraining potentials were applied $\frac{V_1(r)}{T^*} = \begin{cases} r, & x < R_c \\ 0, & else \end{cases}$ and $\frac{V_2(r)}{T^*} = \begin{cases} r, & x \geq R_c \\ 0, & else \end{cases}$, where $V_1$ is applied to Crowder chains only, and $V_2$ is applied to all other components. $r$ corresponds to the Euclidean distance from the center of the lattice. The constraining radius, $R_c = \sqrt[3]{\frac{N_{molecules} - N_{crowder}}{\pi}}^{\frac{1}{3}}$, is such that the crowder-free bead density within the bounding sphere is $\rho_0 \approx 0.75$. $N_{molecules}$ and $N_{crowder}$ correspond to the total number of beads corresponding to the interacting molecules and crowder, respectively. The constraining potential causes the non-crowder chains to be localized to the center of the lattice. The temperature is then discontinuously changed to $3.2T^*$. The system is annealed, and temperature is linearly reduced to $1.2T^*$ over $1 \times 10^8$ steps. The initial constraining potential is then turned off. The temperature is discontinuously changed to $1.1T^*$, followed by $1 \times 10^8$ steps. Again, the temperature is discontinuously reduced to $1T^*$ and $2.11 \times 10^{10}$ steps are performed.

Data are acquired in the last $5 \times 10^9$ steps at a frequency of $2.5 \times 10^6$ steps. The averages from each replicate of the 5 replicates is used to generate Fig. 5C.

Table 3 shows the exact number of molecules used for each of the mixtures. A ratio of 1 to 5 chains is used for DNA-Protein mixtures, 1 to 2 chains for DNA-RNA mixtures, and 2 to 5 chains for RNA-Protein mixtures. This is to keep the number of modules from each species the same in the simulations.

**Table 3: Molecule numbers for simulations of binary mixtures**

| | DNA | TFAM | TFB2M | POLRMT | RNA | Crowder |
|---|---|---|---|---|---|---|
| **DNA+TFAM** | 1000 | 5000 | 0 | 0 | 0 | 5000 |
| **DNA+TFB2M** | 1000 | 0 | 5000 | 0 | 0 | 5000 |
| **DNA+POLRMT** | 1000 | 0 | 0 | 5000 | 0 | 5000 |
| **TFAM+TFB2M** | 0 | 5000 | 5000 | 0 | 0 | 5000 |
| **TFAM+POLRMT** | 0 | 5000 | 0 | 5000 | 0 | 5000 |
| **TFB2M+POLRMT** | 0 | 0 | 5000 | 5000 | 0 | 5000 |
| **RNA+DNA** | 1000 | 0 | 0 | 0 | 2000 | 5000 |
| **RNA+TFAM** | 0 | 5000 | 0 | 0 | 2000 | 5000 |
| **RNA+TFB2M** | 0 | 0 | 5000 | 0 | 2000 | 5000 |
| **RNA+POLRMT** | 0 | 0 | 0 | 5000 | 2000 | 5000 |

***Full Mixtures***

For the simulations of the full set of components including RNA, the following protocol was used. For $5 \times 10^7$ steps, the simulation temperature was raised to $1000T^*$, all interactions were ignored, and the constraining potentials $\frac{V_3(r)}{T^*} = \begin{cases} r, & x < R' \\ 0, & else \end{cases}$ and $\frac{V_4(r)}{T^*} = \begin{cases} r, & x \geq R' \\ 0, & else \end{cases}$ were used. $V_3$ is applied to Crowder, DNA and RNA chains, and $V_4$ is applied to TFAM, TFB2M, and POLRMT. The constraining radius $R' = \sqrt[\frac{1}{3}]{\frac{N_{proteins}}{\pi}}$, where $N_{proteins}$ is the total number of beads for the proteins, is such that the bead density inside the bounding sphere is approx. 0.75. This results in TFAM, TFB2M and POLRMT localizing towards the center of the lattice. The temperature was then discontinuously reduced to $4T^*$, and the interactions were turned on. The constraining potentials $V_1(r)$ and $V_2(r)$, from above, were then applied. $1 \times 10^8$ steps were performed, the system was annealed, and the temperature is linearly reduced from $4T^*$ to $2T^*$. This results in all non-crowder chains to be localized near the center of the lattice. The temperature was then discontinuously changed to $1.5T^*$ and $1 \times 10^8$ steps were performed, followed by one more discontinuous temperature change to $1T^*$. $4.61 \times 10^{10}$ steps were performed and data were acquired at a frequency of $2.5 \times 10^6$ steps over the last $1 \times 10^{10}$ steps, and the average was stored. The average over the 5 replicates was used to generate Fig. 5C, Fig. S7.

As in the binary mixture simulations, the DNA to protein ratio is kept at 1 to 5. To assess the effects of RNA, different amounts of RNA chains were included in the systems while all other components were kept fixed. RNA chains were added in increments of 100 until there were 1,000 chains, or as many RNA chains as there were DNA chains in the system. The numbers are in Table 4 below.

**Table 4: Molecule numbers for simulations of full set of components**

| Time | DNA | TFAM | TFB2M | POLRMT | RNA | Crowder |
|---|---|---|---|---|---|---|
| $t_0$ | 1000 | 5000 | 5000 | 5000 | 1 | 5000 |
| $t_1$ | 1000 | 5000 | 5000 | 5000 | 100 | 5000 |
| $t_2$ | 1000 | 5000 | 5000 | 5000 | 200 | 5000 |
| $t_3$ | 1000 | 5000 | 5000 | 5000 | 300 | 5000 |
| $t_4$ | 1000 | 5000 | 5000 | 5000 | 400 | 5000 |
| $t_5$ | 1000 | 5000 | 5000 | 5000 | 500 | 5000 |
| $t_6$ | 1000 | 5000 | 5000 | 5000 | 600 | 5000 |
| $t_7$ | 1000 | 5000 | 5000 | 5000 | 700 | 5000 |
| $t_8$ | 1000 | 5000 | 5000 | 5000 | 800 | 5000 |
| $t_9$ | 1000 | 5000 | 5000 | 5000 | 900 | 5000 |
| $t_{10}$ | 1000 | 5000 | 5000 | 5000 | 1000 | 5000 |

To generate the density profiles corresponding to the components, the system center-of-mass (COM) is used as the reference point. From this reference point, radial number histograms of each component are generated. These number histograms are then normalized to generate the radial density profiles. For more details, see the supplementary index in the work of Ruff et al., (*63*).

**Statistical methods**

All imaging experiments had at least three fields of view and were independently repeated at least two-four times, as indicated. Data were pooled together from all experimental replicates, with mean and error reported as indicated in the figure legends.

**Figure S1**

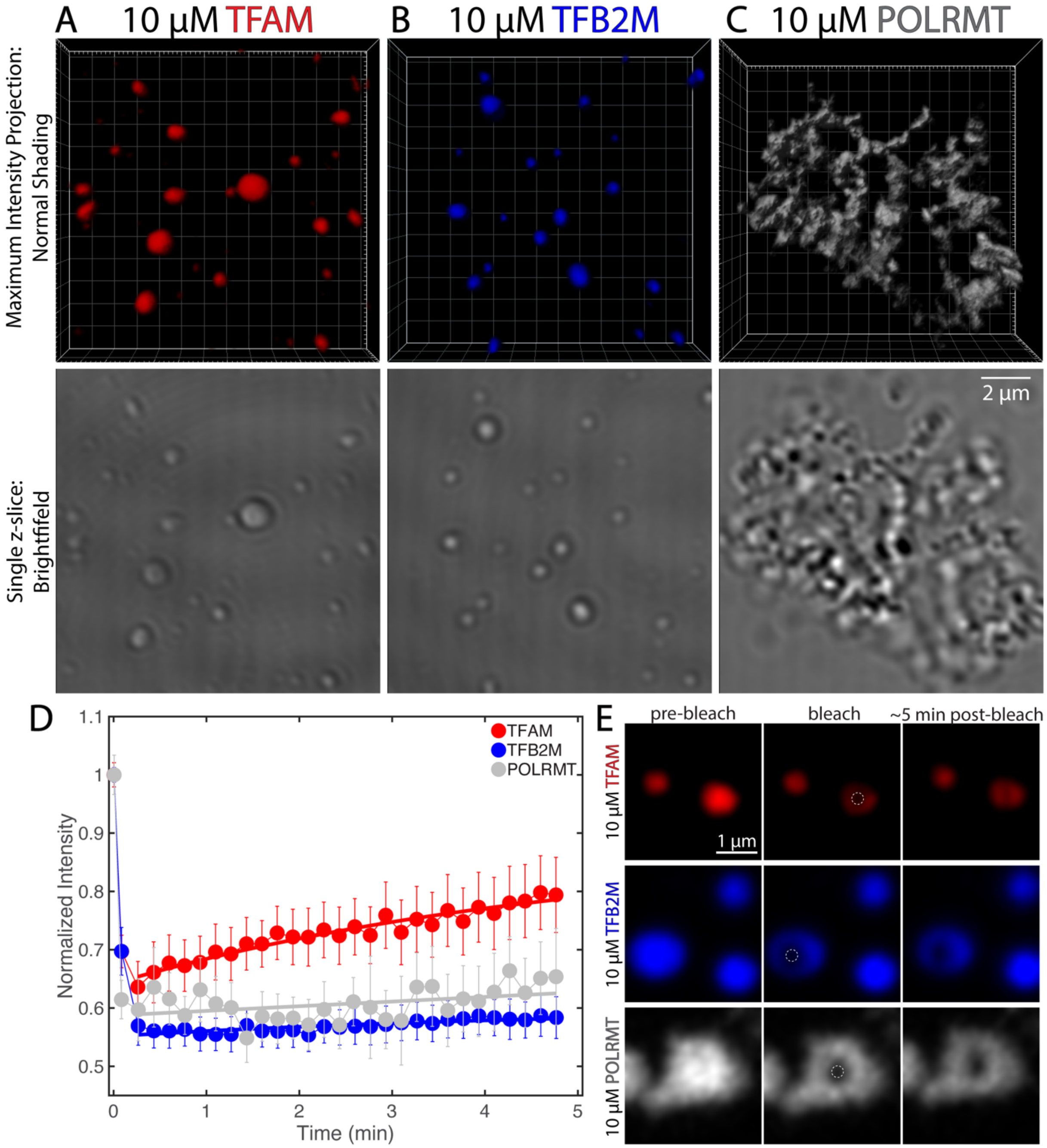


**Fig. S1** Laser-scanning confocal microscopy images of 10 μM TFAM (A), 10 μM TFB2M (B), and 10 μM POLRMT (C) with trace DyLight labelling (see Methods) in the presence of 10% PEG. Top row is a volume projection using normal shading (Imaris) and bottom row is a brightfield image of a single z-slice. Scale bar = 2 μm. (D) Recovery after photobleaching a ~1 μm sized spot for 10 μM TFAM (red), 10 μM TFB2M (blue), or 10 μM POLRMT (grey) droplets in the presence of 10% PEG. Droplets were imaged approximately ~30 minutes after mixing. Error bars are s.e.m., where n = ~10 droplets per condition. Note: intensities were normalized relative to the total intensity of the droplet. (E) Images of representative drops before photobleaching, immediately after photobleaching, and approximately five minutes post-photobleaching for TFAM (red), TFB2M (blue), and POLRMT (grey). Scale bar = 1 μm.

**Figure S2**

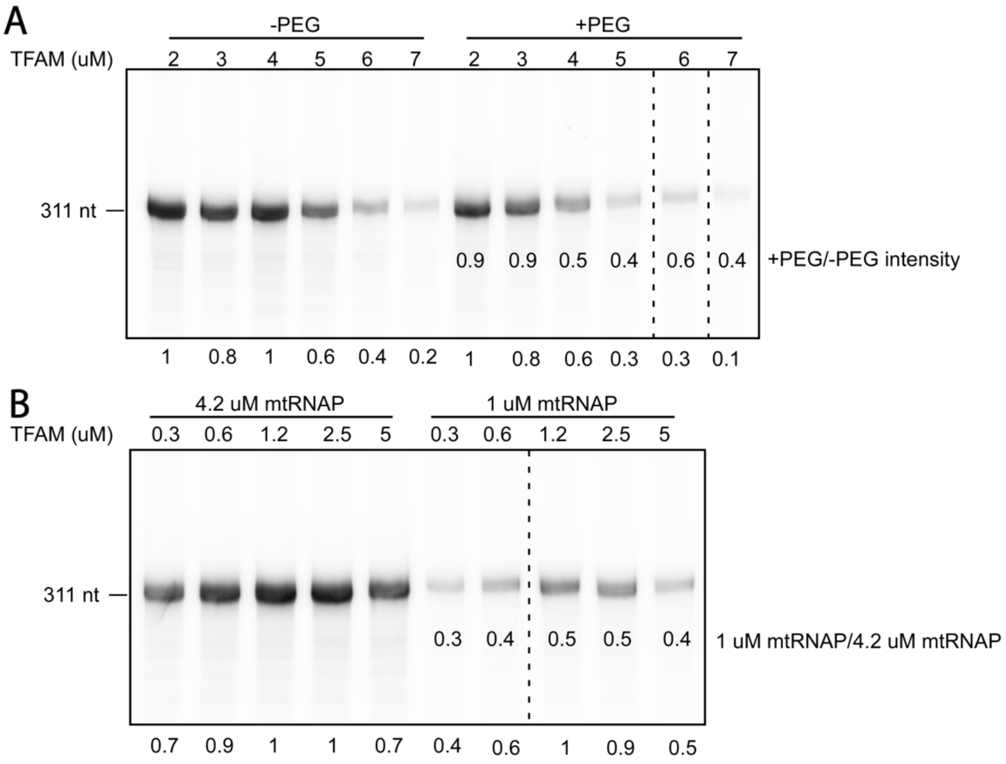


**Fig. S2** (A) RNA blot of transcription run-off assays showing titration of TFAM in presence (+) and absence (–) of 5% PEG using 7X concentrations: 0.35 μM DNA, 4.2 μM POLRMT, 4.2 μM TFB2M and 2-7 μM TFAM (as indicated) in transcription buffer (20 mm Tris HCl, pH 7.9, 10 mM MgCl2, 20 mM $\beta$ME, 2 mM NTP (each), trace P32-GTP, and ~80 mM NaCl. Total RNA product relative to 2 μM TFAM –PEG are reported below the blot. Ratios of +/– PEG are reported underneath bands. (B) RNA blot of transcription run-off assays showing titration of TFAM in absence (–) of PEG for different concentrations of POLRMT using 7X concentrations: 0.35 μM DNA, 1 or 4.2 μM POLRMT, 4.2 μM TFB2M and 0.3-5 μM TFAM (as indicated) in transcription buffer (20 mm Tris HCl, pH 7.9, 10 mM MgCl2, 20 mM $\beta$ME, 2 mM NTP (each), trace P32-GTP, and ~70-80 mM NaCl. Total RNA product relative to 1.2 μM TFAM for each concentration of POLRMT are reported below the blot. Ratios of 1 μM/4.2 μM POLRMT are reported underneath bands.

**Figure S3**

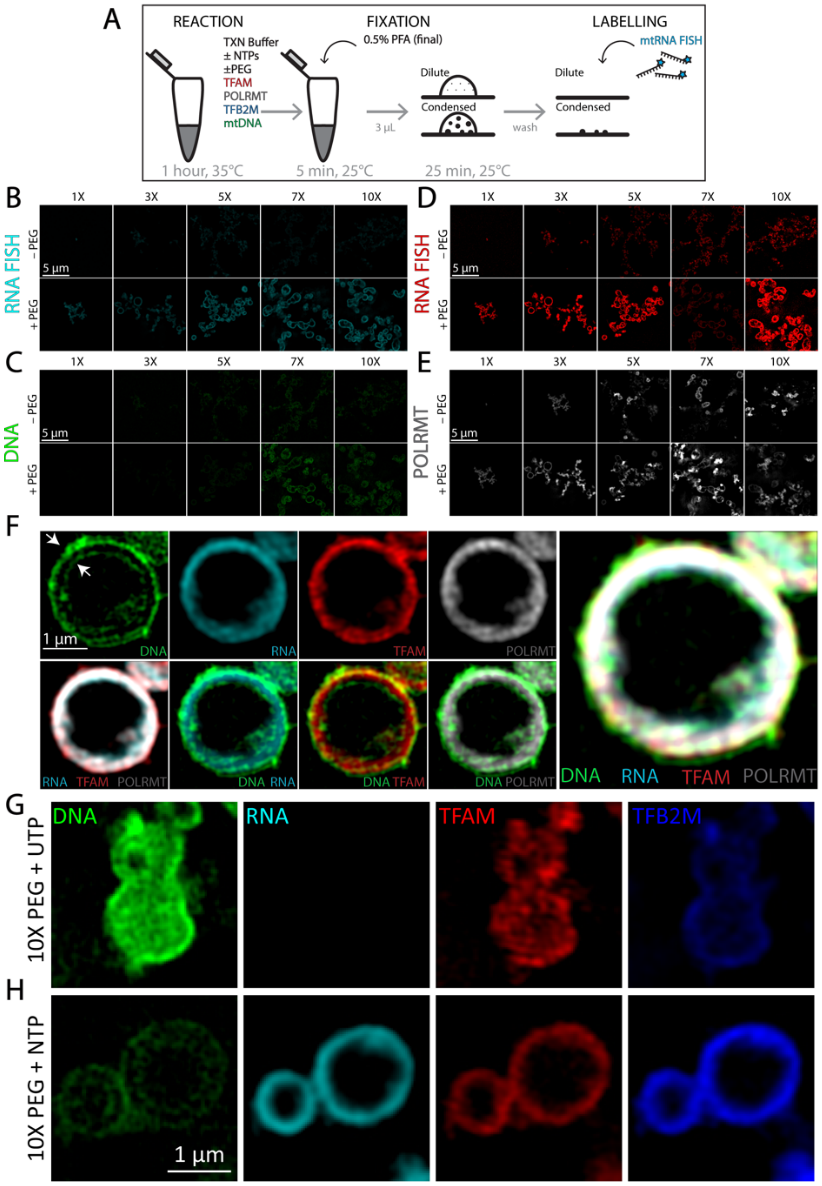


**Fig. S3** (A) Schematic diagram of transcription reactions at 35C for 1 hour followed by fixation on glass coverslips and RNA FISH labelling (see Methods). (B-E) SIM images of reactions after RNA FISH labelling for individual channels: RNA (B), DNA (C), TFAM (D), and POLRMT (E). Top row is without PEG (–PEG) and bottom row is with PEG (5%, + PEG) for 1X-10X conditions. Scale bar = 5 µm for all images. Contrast settings are equivalent for images in the same channel. (F) Images of a large vesicle after 1 hour of transcription (7X, 5% PEG, NTPs). Panels include single channel of DNA, RNA, TFAM, and POLRMT (top row), overlays of RNA/TFAM/POLRMT, DNA/RNA, DNA/TFAM, and DNA/POLRMT (bottom row), and a four-channel overlay (right). Arrows indicate the outer an inner lining of DNA. Scale bar =1 µm. (G,H) Images of condensates after 1 hour of incubation (10X, 5% PEG) with UTP (G) or NTP (H). Panels include single channel of DNA (green), RNA (cyan), TFAM (red), and TFB2M (blue). Scale bar =1 µm.

### Figure S4

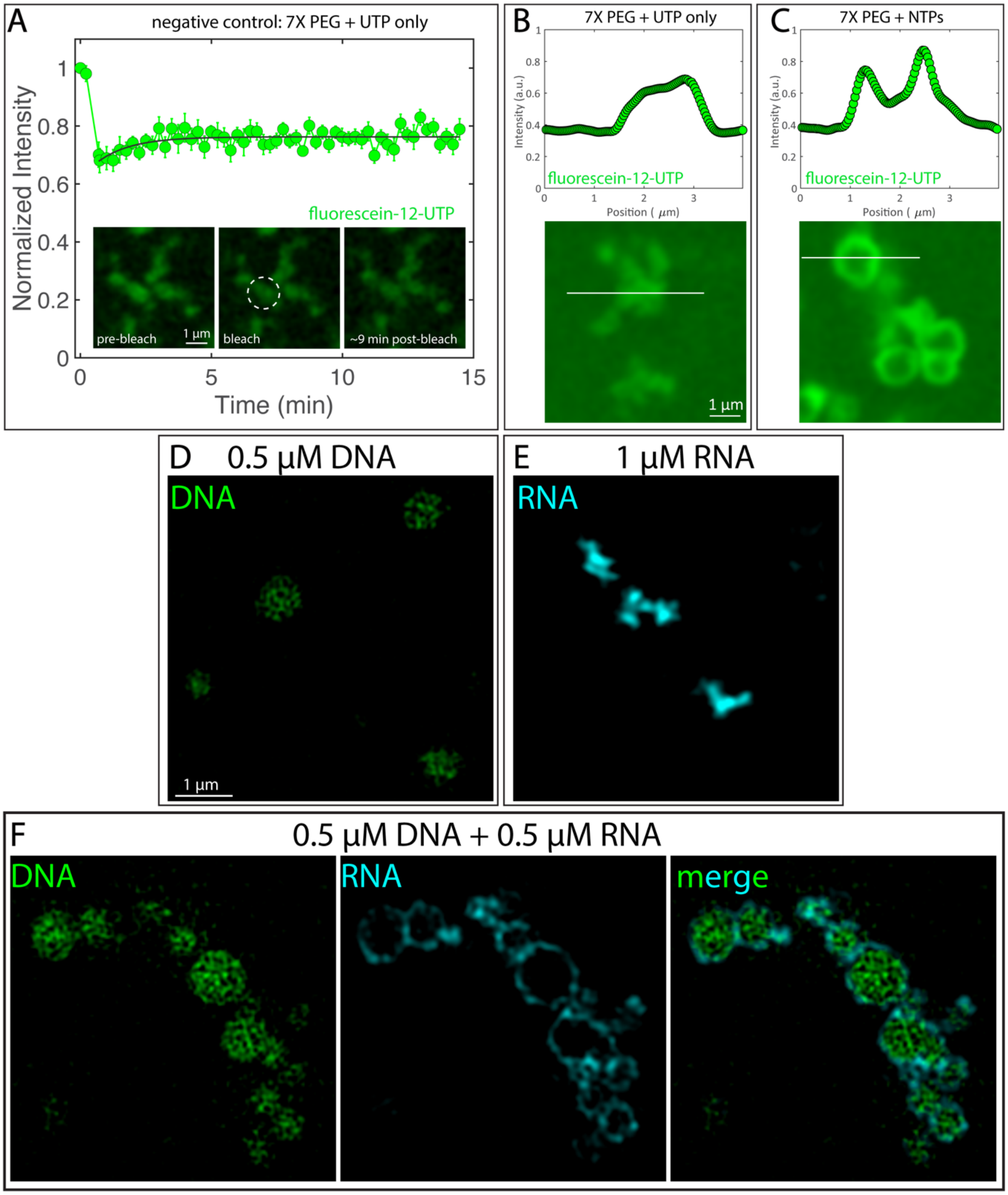


**Fig. S4** (A) FRAP recovery for non-reacting droplets (8 mM UTP) for 7X and 5% PEG conditions. Inset shows condensates pre-bleach, bleach, and 9 min post-bleach. Dashed circle represents region that was bleached. Scale bar = 1 µm. n = 10 droplets and error bars = standard error of the mean. (B,C) Line profiles of fluorescein-12-UTP localization in condensates formed at 7X +PEG conditions in the presence of UTP (B, negative control) or NTP (C, transcription). Intensities are matched between the two images and reflect the background levels of UTP as well as enrichment within the condensate (B) or at the periphery of vesicles (C). Scale bar = 1 μm. (D-F) SIM images of i*n vitro* assays of pure DNA (0.5 μM, DAPI, green) (D), pure RNA (1 μM, CF-640R-UTP, cyan) (E), or DNA+RNA (0.5 μM, each). Buffer was 20 mM Tris-HCl, pH ~7.9, 10 mM $MgCl_2$, 20 mM BME, 80 mM NaCl, and 10% PEG. Solutions were made by preparing buffer, followed by addition of DNA only (D), RNA only (E), or DNA followed by RNA (F). 3 μl of mixture were applied to a pluronic treated coverslip and sealed with a silicone well (Grace Bio-Labs) and imaged 30-60 minutes post mixing, all at room-temperature. Scale bar = 1 μm.

**Figure S5**

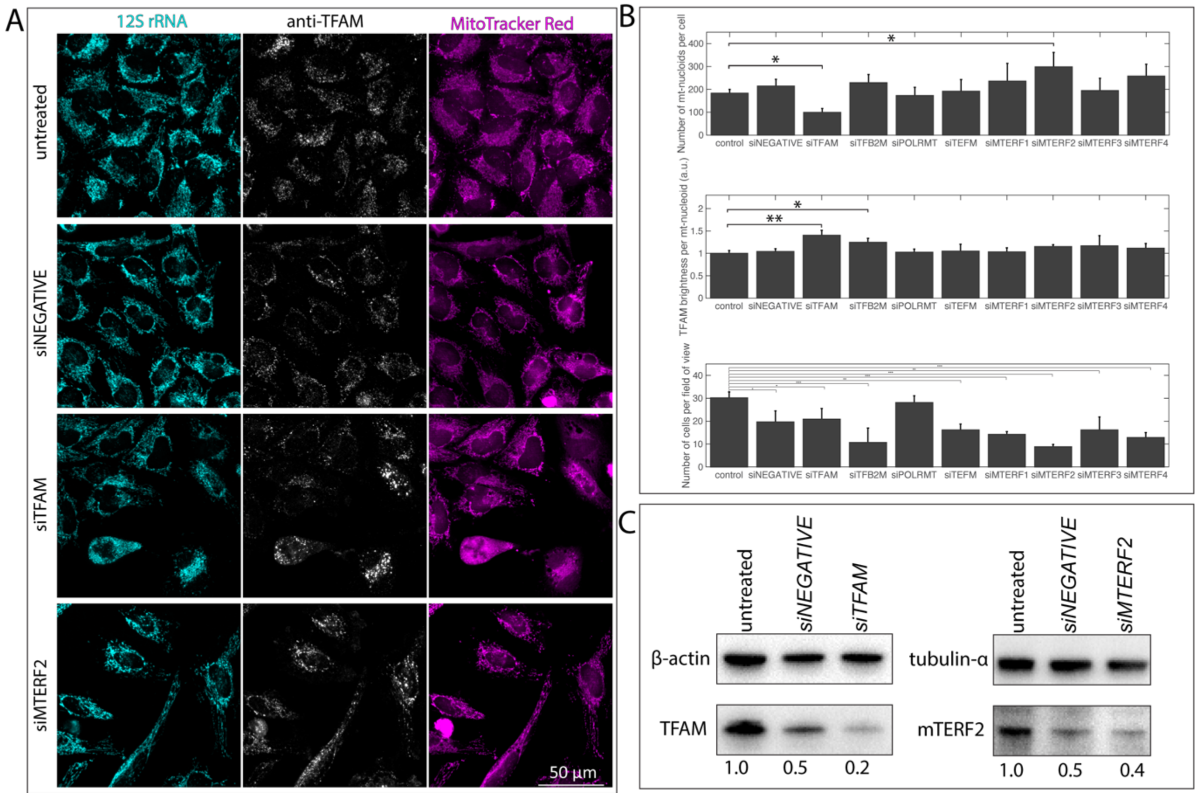


**Fig. S5** (A) Maximum intensity images from high-throughput confocal imaging after siRNA knockdown of most significant conditions, where left panels are of 12S rRNA (RNA FISH, cyan), middle panels are of TFAM (anti-TFAM, gray scale), and right panels are of mitochondria (MitoTracker Red, magenta), for untreated, siNEGATIVE, siTFAM, and siMTERF2. Images shown represent approximately 25% of entire field of view as detected by camera on CV7000 Yokogawa high-throughput cytological discovery system. Scale bar = 50 µm. (B) Quantitative analysis of high-throughput imaging after siRNA treatment, including number of mt-nucleoids per cell, TFAM brightness per mt-nucleoid, and number of cells. Analysis is of one experimental replicate with three technical replicates, each with five fields of view, where error bars = standard deviation. P-value for ANOVA test was $P < 0.01$ for number mt-nucleoids per cell, $P < 0.01$ for TFAM brightness per mt-nucleoid, and $P < 0.001$ for cell number. For individual pairs, a least significant difference procedure was performed, where *$P<0.05$, **$P<0.01$, and ***$P<0.001$. (C) Left: Western blot confirming TFAM (anti-TFAM) depletion for untreated, siNEGATIVE, and siTFAM in HeLa cells after 72 hours. Loading control was actin (anti-beta-actin). Right: Western blot confirming mTERF2 (anti-mTERF2) depletion for untreated, siNEGATIVE, and siMTERF2 in HeLa cells after 72 hours. Loading control was tubulin (anti-tubulin-alpha).

### Figure S6

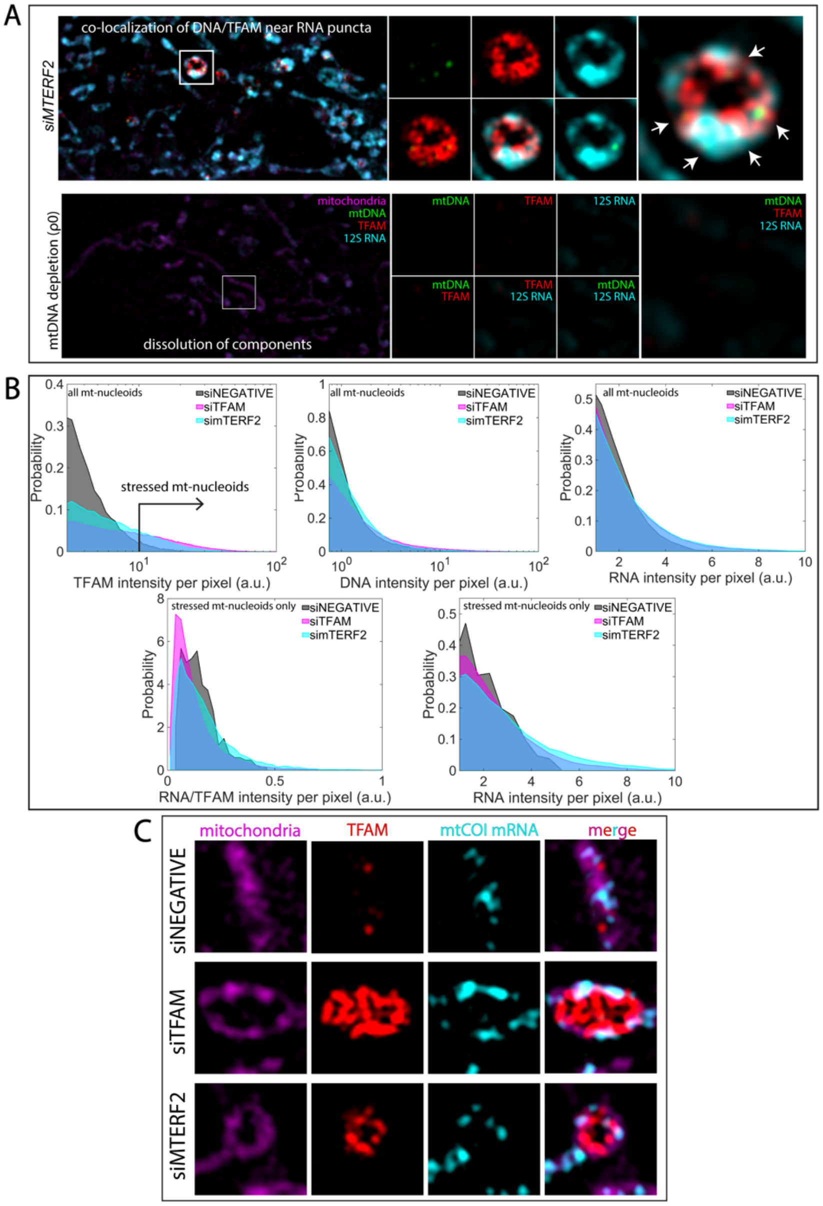


**Fig. S6** (A) SIM images of mitochondrial components after 72 hours of siMTERF2 (top) or mtDNA depletion (bottom). Leftmost panels are the merged image of the zoomed-out version of the mitochondrial network, where mitochondria are in magenta (MitoTracker Red), mtDNA is in green (anti-DNA), TFAM is in red (anti-TFAM), and 12S mtRNA is in cyan (RNA FISH). Arrows in (A) indicate mtDNA puncta co-localizing with TFAM, and 12S rRNA. Scale bar = 1 µm. White box indicates region of interest (ROI). Middle panels are signal or two-channel overlays of the ROI. Rightmost panels are the three-channel overlay for the ROI. Scale bar = 0.2 µm. Intensities across each channel are matched the same as in siRNA conditions (Fig. 4). n = 4 independent experimental replicates. (B) Probability distribution of intensity of TFAM, mtDNA, 12S rRNA, or 12S rRNA/TFAM within all segmented nucleoids or stressed nucleoids (see black arrow in Fig. 4), as indicated, where gray = siNEGATIVE, magenta = siTFAM, and cyan = siMTERF2. (C)

SIM images of mitochondria (magenta), TFAM (red), COI mRNA (cyan), and merged channels for siNEGATIVE, siTFAM, and siMTERF2.

**Figure S7**

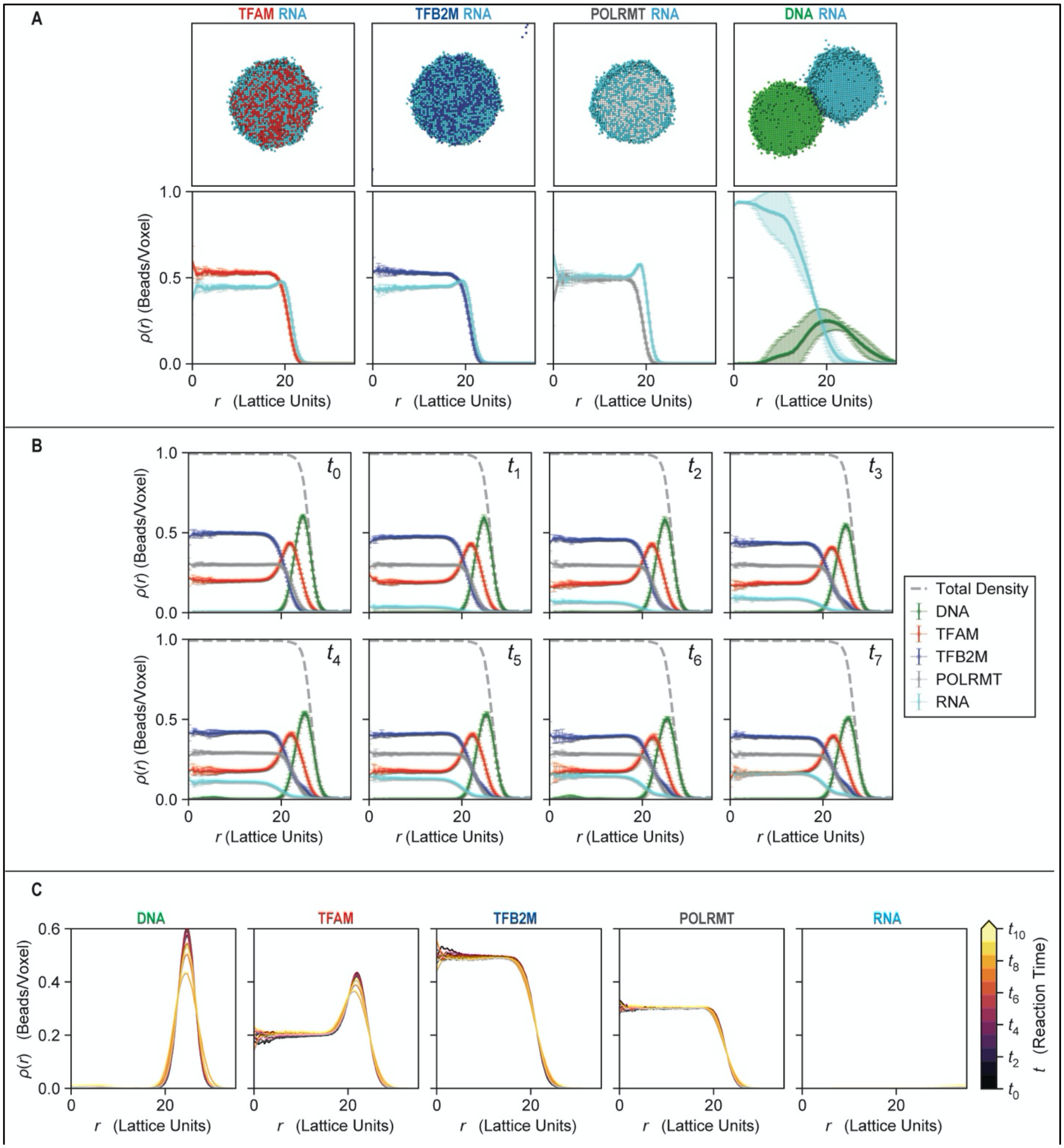


**Fig. S7** (A) Representative snapshots and density profiles for all binary mixtures containing RNA with Model A. For clarity, the Crowder is not shown. RNA has favorable interactions with TFAM, TFB2M and POLRMT and thus generates well mixed droplets in those cases. Since RNA and DNA do not have favorable interactions, RNA and DNA make separate droplets. (B) Density profiles of each component with Model A. Each panel represents a different amount of RNA in the system, or a time point further along the transcription reaction. At the very start when there is no RNA in the system, the components form a multi-phasic layered droplet. POLRMT and TFB2M are mostly in the interior. TFAM is found in the interior but also forms a higher density layer near the DNA. The DNA is most exterior making a DNA-TFAM shell. As more RNA is introduced, more RNA becomes included towards the interior of the droplet. Since the total density cannot be higher than 1, the density of TFB2M is reduced in the interior and is pushed outwards. This further causes the broadening of the exterior DNA-TFAM layer. (C) Density profiles of each component, in separate panel, with Model B as a function of RNA amount, or reaction time. Since the RNA can favorably interact with the Crowder, we can see that there is no RNA inside the transcriptional condensate. TFB2M and POLRMT, which are mostly towards the interior are unaffected by the increased amount of RNA. The outermost layers which include DNA and TFAM are slightly affected. As more RNA is included, more of the Crowder is engaged in interactions with RNA, which reduces the effective amount of Crowder for the transcriptional components.
